# E-box independent chromatin recruitment turns MYOD into a transcriptional repressor

**DOI:** 10.1101/2024.12.05.627024

**Authors:** Chiara Nicoletti, Jimmy Massenet, Andreas P. Pintado-Urbanc, Leah J. Connor, Monica Nicolau, Swetha Sundar, Mingzhi Xu, Anthony Schmitt, Wenxin Zhang, Zesen Fang, Tsz Ching Indigo Chan, Stephen J. Tapscott, Tom H. Cheung, Matthew D. Simon, Luca Caputo, Pier Lorenzo Puri

## Abstract

MYOD is an E-box sequence-specific basic Helix-Loop-Helix (bHLH) transcriptional activator that, when expressed in non-muscle cells, induces nuclear reprogramming toward skeletal myogenesis by promoting chromatin accessibility at previously silent loci. Here, we report on the identification of a previously unrecognized property of MYOD as repressor of gene expression, via E-box-independent chromatin binding within accessible genomic elements, which invariably leads to reduced chromatin accessibility. MYOD-mediated repression requires the integrity of functional domains previously implicated in MYOD-mediated activation of gene expression. Repression of mitogen-and growth factor-responsive genes occurs through promoter binding and requires a highly conserved domain within the first helix. Repression of cell-of-origin/alternative lineage genes occurs via binding and decommissioning of distal regulatory elements, such as super-enhancers (SE), which requires the N-terminal activation domain as well as two chromatin-remodeling domains and leads to reduced strength of CTCF-mediated chromatin interactions. Surprisingly, MYOD-mediated chromatin compaction and repression of transcription do not associate with reduction of H3K27ac, the conventional histone mark of enhancer or promoter activation, but with reduced levels of the recently discovered histone H4 acetyl-methyl lysine modification (Kacme). These results extend MYOD biological properties beyond the current dogma that restricts MYOD function to a monotone transcriptional activator and reveal a previously unrecognized functional versatility arising from an alternative chromatin recruitment through E-box or non-E-box sequences. The E-box independent repression of gene expression by MYOD might provide a promiscuous mechanism to reduce chromatin accessibility and repress cell-of-origin/alternative lineage and growth factor/mitogen-responsive genes to safeguard the integrity of cell identity during muscle progenitor commitment toward the myogenic lineage.

## Introduction

MYOD has a unique property to activate skeletal myogenesis upon ectopic expression in non-muscle cells – also known as MYOD-mediated trans-differentiation or myogenic conversion of somatic cells^1,2^ – which reflects its function as endogenous activator of skeletal myogenesis in muscle stem cells (MuSCs) during skeletal muscle regeneration^3,4^. This property relies on MYOD ability to bind nucleosomes at previously silent loci in cooperation with pioneer factors, such as Pbx1/Meis^5^, followed by signal-dependent recruitment of histone acetyltransferases and SWI/SNF chromatin remodeling complex^6–16^ to promote chromatin accessibility and enable full recognition and binding to specific E-box sequences (wherein the central di-nucleotide is GC or GG)^17,18^. E-box-driven heterodimerization with E2A gene products (E12 and E47)^19^ enables MYOD to activate transcription of target skeletal muscle-specific genes^20^. Previous studies have identified specific domains that confer on MYOD the property as a DNA sequence-specific transcriptional activator^21^. Within the bHLH region^22^, the basic domain restricts MYOD DNA binding affinity to myogenic E-box motifs^23^ and the HLH domain mediates heterodimerization with E12 or E47^24^. Moreover, an acidic activation domain (AD) located at the N-terminus^25^ and two chromatin remodeling domains (CRDs located at the C/H-rich domain and C-terminus)^26^ cooperate to activate transcription of target genes.

Recent work has extended our knowledge of MYOD-mediated activation of skeletal myogenesis, by revealing its pervasive binding throughout the genome^27^ and its role as organizer of the 3D genome architecture^27–33^. While MYOD-mediated activation of gene expression has been extensively studied since its discovery as a myogenic determination factor, the expression of MYOD in proliferating, undifferentiated muscle progenitors has been puzzling. MYOD expression coincides with a stage in which activation of muscle gene expression has not yet occurred, suggesting that MYOD could exert functions alternative to the activation of muscle genes. Interestingly, in activated MuSCs during skeletal muscle regeneration, the expression of endogenous MYOD coincides with the downregulation of a subset of genes invariably assigned to gene ontology processes related to cell-of-origin, alternative lineages, and growth factor responsive genes^31,32,34^. Likewise, gene expression analysis has revealed specific patterns of gene repression during MYOD-mediated myogenic conversion of non-muscle somatic cells^35,36^. However, while the potential function of MYOD as a transcriptional repressor has been sparsely reported by previous works^37–43^, the potential mechanism that accounts for this putative functional property remains elusive.

Here, we report on the identification of MYOD as a direct repressor of gene expression through binding to non-E-box motifs. Functional domains of MYOD implicated in the activation of gene expression (*i.e.,* the N-terminal AD, the two CRDs and the first helix) are also required for the repression of specific subsets of genes, by reducing chromatin accessibility and levels of the novel mark of transcriptional activation, acetyl-methyllysine, (Kacme)^44^, at promoters and enhancers of MYOD-repressed genes.

## Results

### MYOD reduces chromatin accessibility at promoters of mitogen-and growth factor-responsive genes during human fibroblast trans-differentiation into skeletal muscle cells

MYOD-mediated trans-differentiation of somatic cells into skeletal muscles^1,2^ provides an optimal experimental platform to investigate the molecular, genetic and epigenetic mechanism by which MYOD coordinates nuclear reprogramming of non-muscle cells toward the myogenic lineage. In previous studies we have exploited this model to demonstrate that, upon inducible expression in IMR90 human fibroblasts, MYOD re-organizes the 3D genome architecture by rewiring high-order chromatin interactions implicated in the formation of boundaries of functional nuclear domains, such as the insulated neighborhoods (INs), within topologically associating domains (TADs)^28^. This process is well appreciated upon the exposure to differentiation cues (differentiation medium - DM), in which cells uniformly undergo terminal differentiation. A more dynamic genome reprogramming occurs during the proliferation of MYOD-expressing IMR90 (IMR90-MYOD) cells, when they are cultured in high serum (growth medium - GM). Indeed, this is an intermediate stage of commitment toward the myogenic lineage that entails the erasure of the previous cell of origin lineage, prior to the activation of the differentiation program. During this transition, culturing cells in high serum-containing growth factors and mitogens mimics the exposure of muscle progenitors to developmental or regeneration cues, which might activate multiple responses and cell lineages if not properly filtered/interpreted. Thus, we sought to focus our analysis on IMR90 cells cultured in high mitogen/growth factor-containing serum (Ext. Fig. 1A). Under these conditions, doxycycline (doxy)-induced MYOD expression did not activate endogenous MYOD, neither promoted the formation of myosin heavy chain (MYHC)-expressing multinucleated terminally differentiated myotubes (Ext. Fig. 1B). Conversely, MYOD expression coincided with downregulation of genes, including the lung fibroblasts lineage gene *GATA6* (Ext. Fig.1C, D and E), the pro-inflammatory cytokine *interleukin 6 (IL6)*, the growth factor-responsive *cFos,* and the extracellular matrix (ECM) component *Fibronectin 1* (*FN1*) (Ext. Fig. 1D), while typical early MYOD-induced differentiation genes, such as *Integrin alpha 7* (*ITGA7)* and *Troponin T2* (*TNNT2)*, were weakly activated (Ext. Fig. 1D). A late differentiation marker – the embryonic *MYH3* – was not induced at this stage (Ext. Fig. 1D). Consistently, RNA-seq analysis of IMR90-MYOD cells revealed that a large proportion (about 1/3) of differentially expressed genes (DEGs) were downregulated, as compared with control IMR90 cells (Fig. 1A and B).

**Fig. 1.**
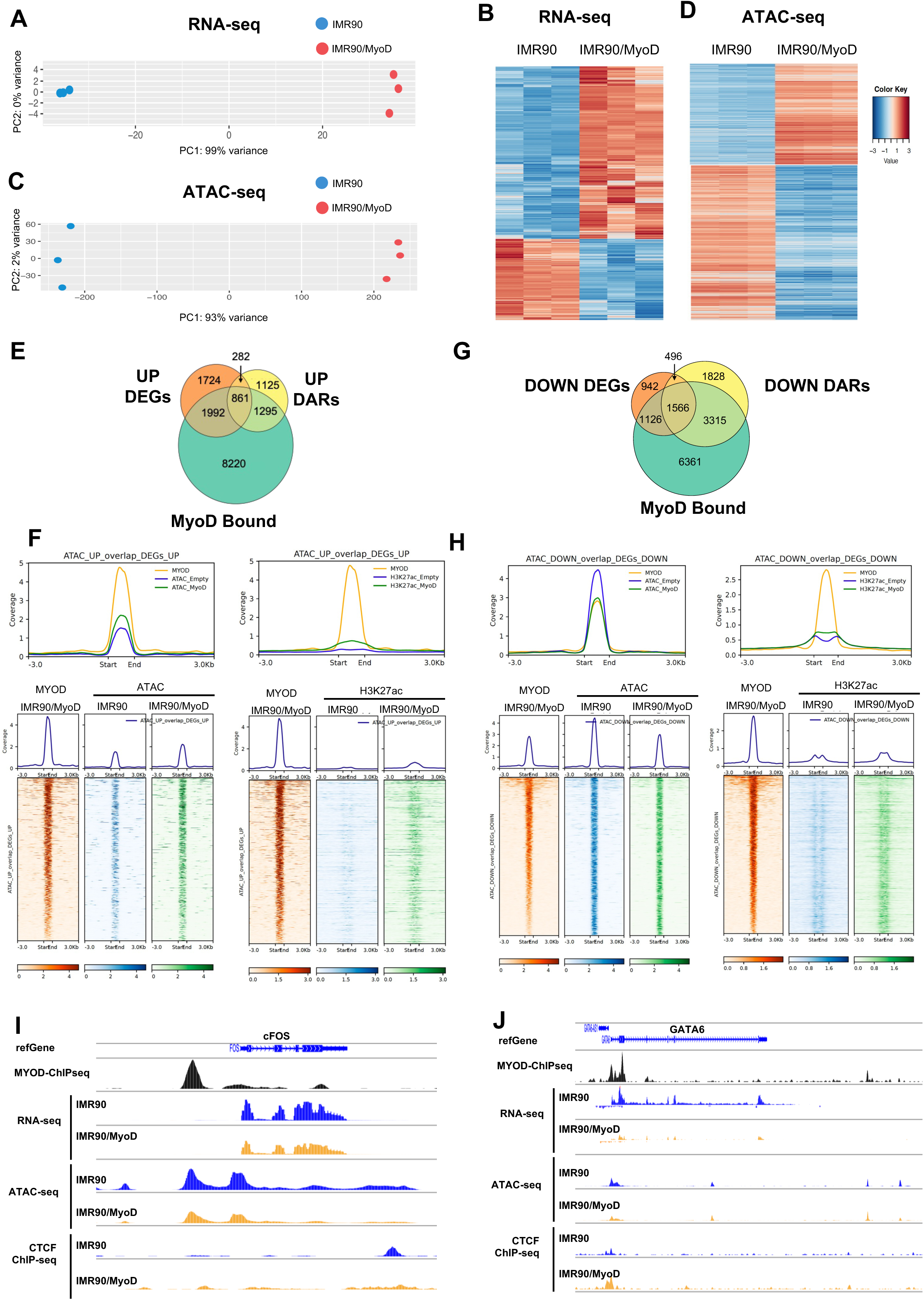
MYOD reduces chromatin accessibility at promoters of mitogen-and growth factor-responsive genes during human fibroblast trans-differentiation into skeletal muscle cells. a) Principal Component Analysis (PCA) of gene expression data for IMR90 and IMR90/MyoD samples. b) Heatmaps of differential gene expression (false-discovery rate (FDR)-adjusted P < 0.05) of IMR90/MyoD vs IMR90. c) Principal Component Analysis (PCA) of chromatin accessibility data for IMR90 and IMR90/MyoD samples. d) Heatmaps of differential promoter accessibility (FDR-adjusted P < 0.05, |logFC|>1.5) of IMR90/MyoD vs IMR90. e) Venn Diagram of the overlap between gene promoters bound by MyoD, differentially expressed (DEGs) and differentially accessible (DARs) for genes up-regulated in IMR90/MyoD vs IMR90. f) Tornado Plots of MyoD ChIP-seq signal in IMR90/MyoD (orange) and ATAC-seq signal in IMR90 (blue) and IMR90/MyoD (green) for differentially accessible promoters (right panels) and H3K27ac ChIP-seq signal in IMR90 (light blue) and IMR90/MyoD (yellow) of up-regulated DEGs. g) Venn Diagram of the overlap between gene promoters bound by MyoD, differentially expressed (DEGs) and differentially accessible (DARs) for genes down-regulated in IMR90/MyoD vs IMR90. h) Tornado Plots of MyoD ChIP-seq signal in IMR90/MyoD (orange) and ATAC-seq signal in IMR90 (blue) and IMR90/MyoD (green) for differentially accessible promoters (right panels) and H3K27ac ChIP-seq signal in IMR90 (light blue) and IMR90/MyoD (yellow) of down-regulated DEGs. i-j) IGV screenshot of the genomic regions of cFos (j) and GATA6 (k). Tracks from top to bottom: refseq gene, MYOD ChIP-seq, RNA-seq tracks in IMR90 (blue) and IMR90/MYOD (orange), ATAC-seq tracks in IMR90 (blue) and in IMR90/MYOD (orange), CTCF ChIP-seq in IMR90 (blue) and in IMR90/MYOD (orange).

Likewise, ATAC-seq shows analogous patterns of reduced and induced chromatin accessibility at promoters (Fig. 1C and D), with a notable higher number of events of reduced chromatin accessibility (Fig. 1D), which indicates that repression of gene expression occurs through extensive chromatin compaction at promoters of repressed genes (Fig. 1D). To determine a causal relationship between MYOD chromatin binding, changes in gene expression and in chromatin accessibility, we integrated MYOD binding events at promoters (by ChIP-seq) with promoters of DEGs (by RNA-seq) and with the chromatin accessibility patterns identified by ATAC-seq in IMR90 vs IMR90-MYOD cells cultured in GM for 24 hours. We found that about 1/3 of MYOD ChIP peaks coincided with binding to more than half of the promoters of upregulated genes or overlapped with increased promoter chromatin accessibility, with only a minority (861 peaks) associated to increased chromatin accessibility at promoters of upregulated genes (Fig. 1E and F). Gene ontology analysis revealed that MYOD binding to promoters with increased chromatin accessibility coincided with the upregulation of genes implicated in biological processes related to skeletal myogenesis and general features of differentiation (Ext. Fig. 2A). Motif analysis of these events showed an invariable association with typical MYOD targets, the myogenic E-box motif (CAGCTG) (Ext. Fig. 2C). Conversely, more than half of MYOD peaks (6007) detected by ChIP-seq coincided with binding to a large majority of promoters of down-regulated genes or overlapped with reduced promoter chromatin accessibility, with 1566 peaks associated with reduced chromatin accessibility at promoters of down-regulated genes (Fig. 1G and H). Gene ontology analysis of DEGs indicates that MYOD-bound promoters with reduced chromatin accessibility coincided with downregulation in genes implicated in cell proliferation, regulation of S phase, mitosis, and other phases of the cell cycle (Ext. Fig. 2B). Motif analysis of these events showed an invariable association to non-E-box motifs, with enrichment in motifs for transcription factors (TFs) implicated in cell cycle regulation and proliferation of muscle progenitors, such as E2F^45–47^ and NFY^48,49^, or serum/growth factor responsive TFs, such as SP1 and Elk family members^50^ (Ext. Fig. 2D). Interestingly, while the increased chromatin accessibility at MYOD-bound promoters of upregulated genes was associated with increased levels of H3K27ac, as expected (Fig. 1F), the reduced chromatin accessibility at MYOD-bound promoters of down-regulated genes was also associated with increased levels of H3K27ac (Fig. 1H), which appears paradoxical, as this is a conventional mark for promoter (as well as enhancer) activation. Representative tracks for two downregulated genes, cFOS and GATA6, are shown (Fig 1I and J).

**Fig. 2.**
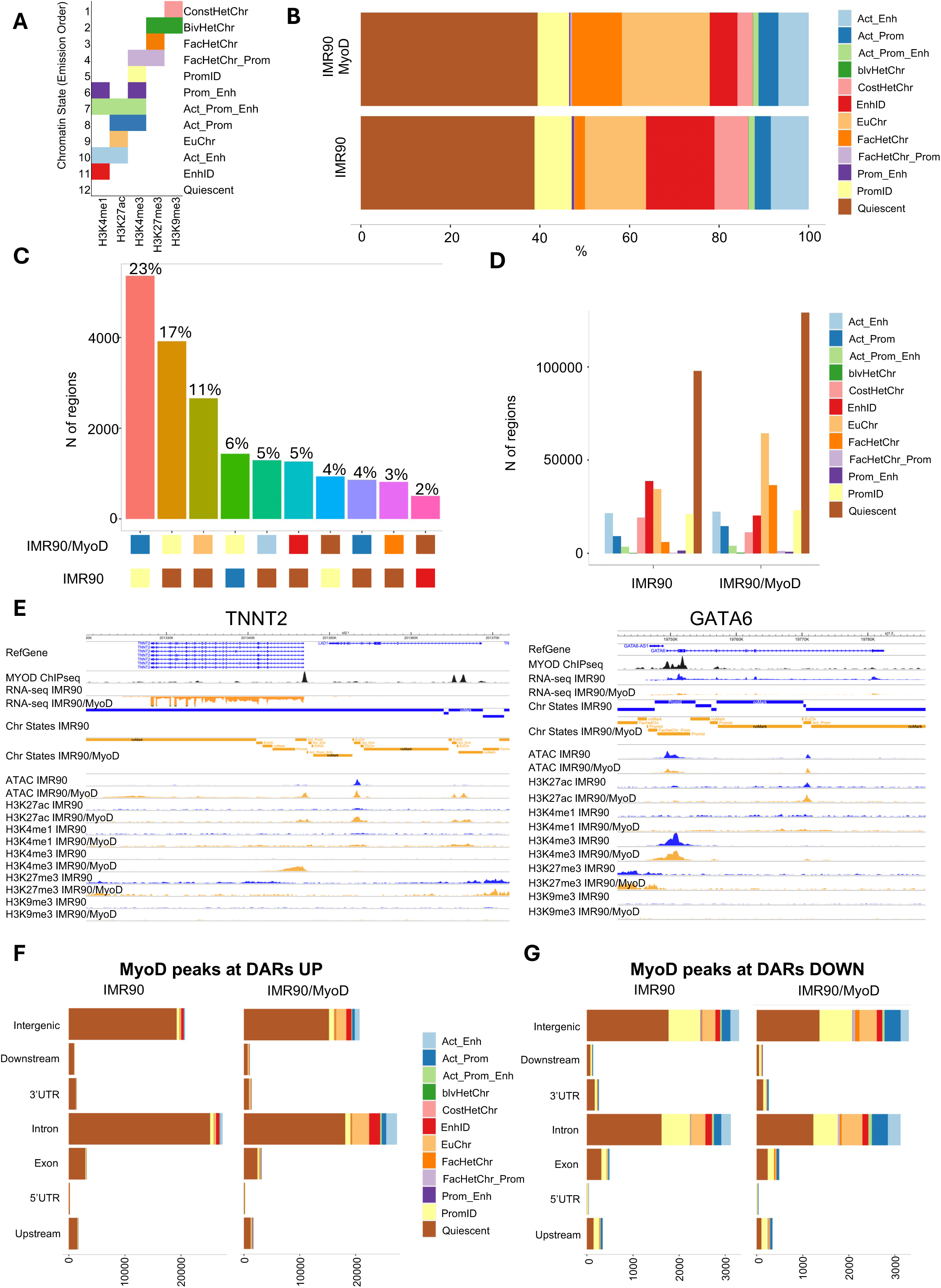
Changes in chromatin states during MYOD-induced IMR90 human fibroblast trans-differentiation into skeletal muscle cells. a) ChromHMM chromatin states emissions based on H3K4me1, H3K27ac, H3K4me3, H3K27me3 and H3K9me3 genome-wide CUT&RUN signal. b) Bar plot of the representation of each chromatin state for the IMR90 and IMR90/MyoD conditions. c) Bar plot of the absolute number of regions for each chromatin state for the IMR90 and IMR90/MyoD conditions. d) Barplot graph number of regions bound by MYOD that change chromatin states between IMR90 and IMR90/MyoD conditions. e) Example IGV tracks with chromatin states at genes activated (*TNNT2*) or repressed (*GATA6*). f-g) Bar plot of absolute number of regions for each chromatin state for ATAC DARs with either increased (f) or decreased (g) accessibility in IMR90 (left panels) and IMR90/MYOD (right panels)

These results suggest that MYOD directly contributes to two distinct programs for genome reprogramming – the activation and repression of different patterns of gene expression - by promoting opposite patterns of chromatin accessibility at promoters of DEGs, via binding to either myogenic E-box or non-E-box motifs.

The different outcome in terms of changes in chromatin accessibility induced by MYOD via chromatin binding at E-box vs non-E-box motifs apparently contradicts the current dogma that MYOD chromatin binding is only driven by the selective affinity for specific myogenic E-box motifs^21–23^. It also challenges the current knowledge on structural and functional properties of MYOD, whereby the presence of two chromatin remodeling domains predicts that, upon recruitment to nucleosomes at target gene loci, MYOD only promotes chromatin accessibility^13–16,26^. We therefore sought to investigate MYOD chromatin binding further within the different outcomes of chromatin accessibility by a deeper analysis, in which we fractioned the MYOD ChIP-seq peaks within differential ATAC-seq peaks into three adjacent genomic windows, at the summit and at the two sides of the peak (Ext. Fig. 3A). Motif analysis revealed again an invariable enrichment of the myogenic E-box at the summit as well as at both sides of peaks of increased chromatin accessibility that coincided with MYOD-bound promoters of activated genes (Ext. Fig. 3B). In contrast, no E-box motifs were detected at the summit of peaks of reduced chromatin accessibility that coincided with MYOD-bound Differentially Accessible Regions (DARs) promoters (Ext. Fig. 3C), with low frequency of non-myogenic E-box motifs detected at the sides.

**Fig. 3.**
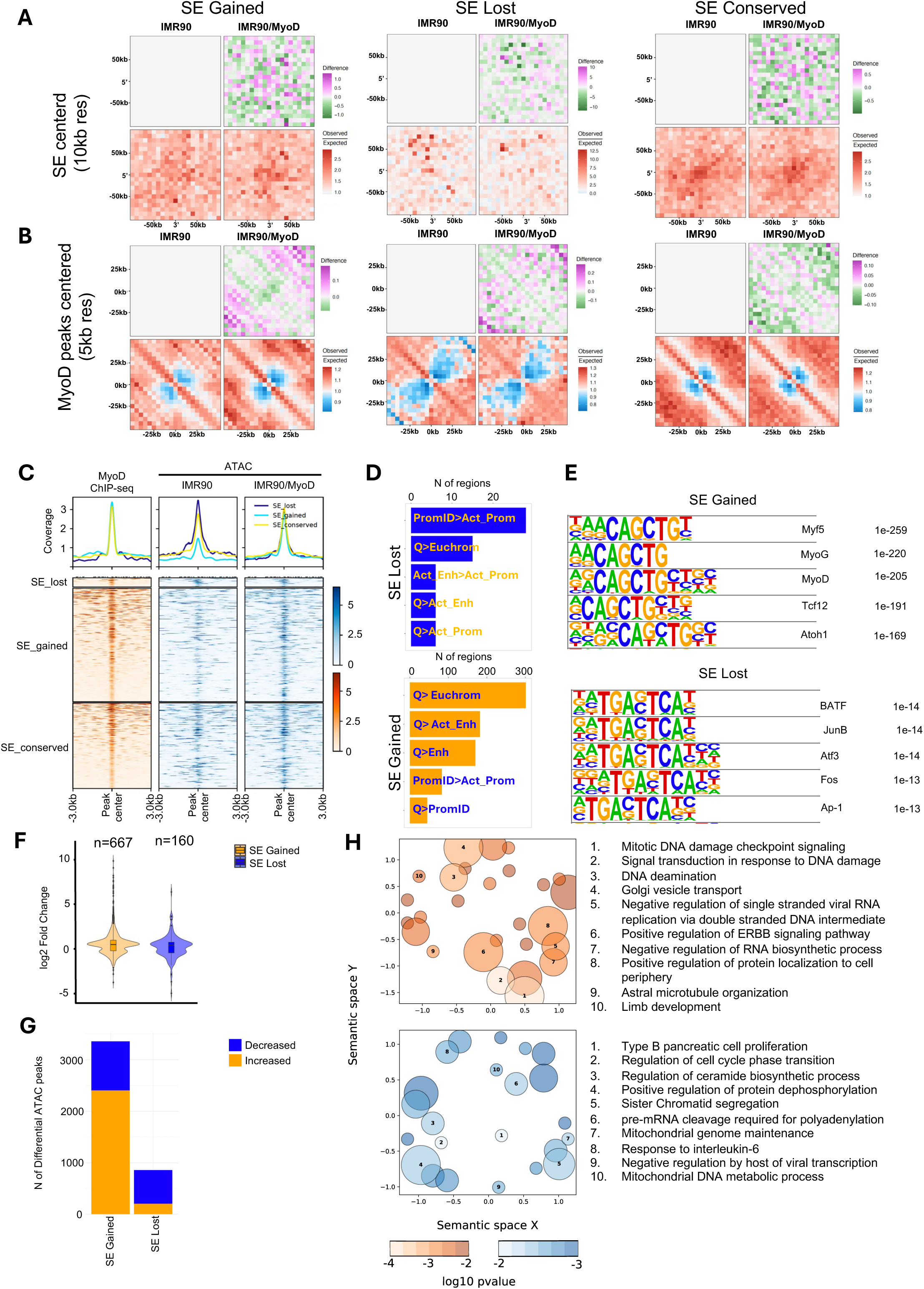
MYOD binds and decommissions SEs of cell-of-origin and alternative lineage genes during human fibroblast trans-differentiation into skeletal muscle cells. a-b) Heatmaps of Aggregate Regions Analysis (ARA) of Hi-C signal at gained (left), lost (center) or conserved (right) super-enhancers bound by MyoD in IMR90/MyoD vs IMR90, using the super-enhancers (a) or MyoD peaks (b) as viewpoint. c) Tornado and aggregate signal plots of MyoD ChIP-seq (orange) and ATAC-seq (blue, center in IMR90, right in IMR90/MyoD) signal at lost (left), gained (center) or conserved (right) super-enhancers bound by MyoD in IMR90/MyoD vs IMR90. d) Top 5 chromatin state changes for SE lost (blue) or gained (orange) e) Top 5 enriched motifs at MyoD peaks bound at lost (top) or gained (bottom) SEs. f) Box plots of log fold change of differentially expressed genes whose promoters interact with gained (orange) or lost (blue) super-enhancers through Hi-C loops. g) Bar plot of differential ATAC peaks overlapping SEs either gained or lost. h) Gene Ontology of biological processes of up-and down-regulated DEGs in (f).

These data reveal unexpected patterns of MYOD binding to promoters of DEGs, leading to opposite patterns of chromatin accessibility associated with either activation or repression of gene expression. While MYOD-mediated activation from promoters entails binding to E-box motifs and leads to induction of genes implicated in early/general features of differentiation, MYOD-mediated repression from promoters occurs through binding to non-E-box motifs and relates to repression of mitogen and growth factor responsive genes.

### Changes in chromatin states during IMR90 human fibroblast trans-differentiation into skeletal muscle cells

As MYOD-mediated repression from promoters did not associate with reduced levels of H3K27ac, we wondered whether other changes in chromatin marks could discriminate the different patterns of MYOD chromatin binding. We therefore performed CUT&RUN analysis to profile the changes in histone marks predictive for promoter (H3K4me3) or enhancer (H3K4me1) identity, for formation of facultative (H3K27me3) or constitutive (H3K9me3) heterochromatin, in addition to H3K27ac, which is a common mark for enhancer and promoter activation. Combinatorial analysis of these marks in IMR90 vs IMR90-MYOD cells identified 12 chromatin states (Fig. 2A), whose dynamics revealed few major features of chromatin state transition, including reduction in constitutive heterochromatin, increased formation of facultative heterochromatin and euchromatin, and partial loss of enhancer identity (Fig. 2B and C). Importantly, when integrated with MYOD ChIP-seq analysis performed in IMR90-MYOD cultured in GM, CUT&RUN analysis of histone modifications revealed that over 2/3 of MYOD-bound genomic elements switch their chromatin state (Fig. 2D and E), again indicating a causal relationship between MYOD chromatin binding at regulatory elements of the genome and changes in chromatin states and conformation. Moreover, integration of CUT&RUN analysis of histone modifications with ATAC-seq datasets revealed a distribution of peaks of increased chromatin accessibility coinciding with increased marks of euchromatin and promoter or enhancer identity across the genome, with a special enrichment at intronic and intergenic elements (Fig. 2F). Conversely, peaks of decreased chromatin accessibility coinciding with increased marks of facultative heterochromatin were observed at promoter and non-promoter regions, distal elements, and were again enriched at intronic and intergenic elements (Fig. 2G).

Further integration of CUT&RUN of histone modifications, MYOD ChIP-seq and ATAC-seq datasets revealed two distinct patterns of changes in chromatin states at promoters of DEGs. While a moderate enrichment in promoter identity (H3K4me3) and activation (H3K27ac) was observed at MYOD-bound promoters of upregulated genes, in association with increased chromatin accessibility (Ext Fig. 4A), the reduction in chromatin accessibility at MYOD-bound promoters of downregulated genes did not coincide with any appreciable changes in histone marks/chromatin states (Ext Fig. 4B).

**Fig. 4.**
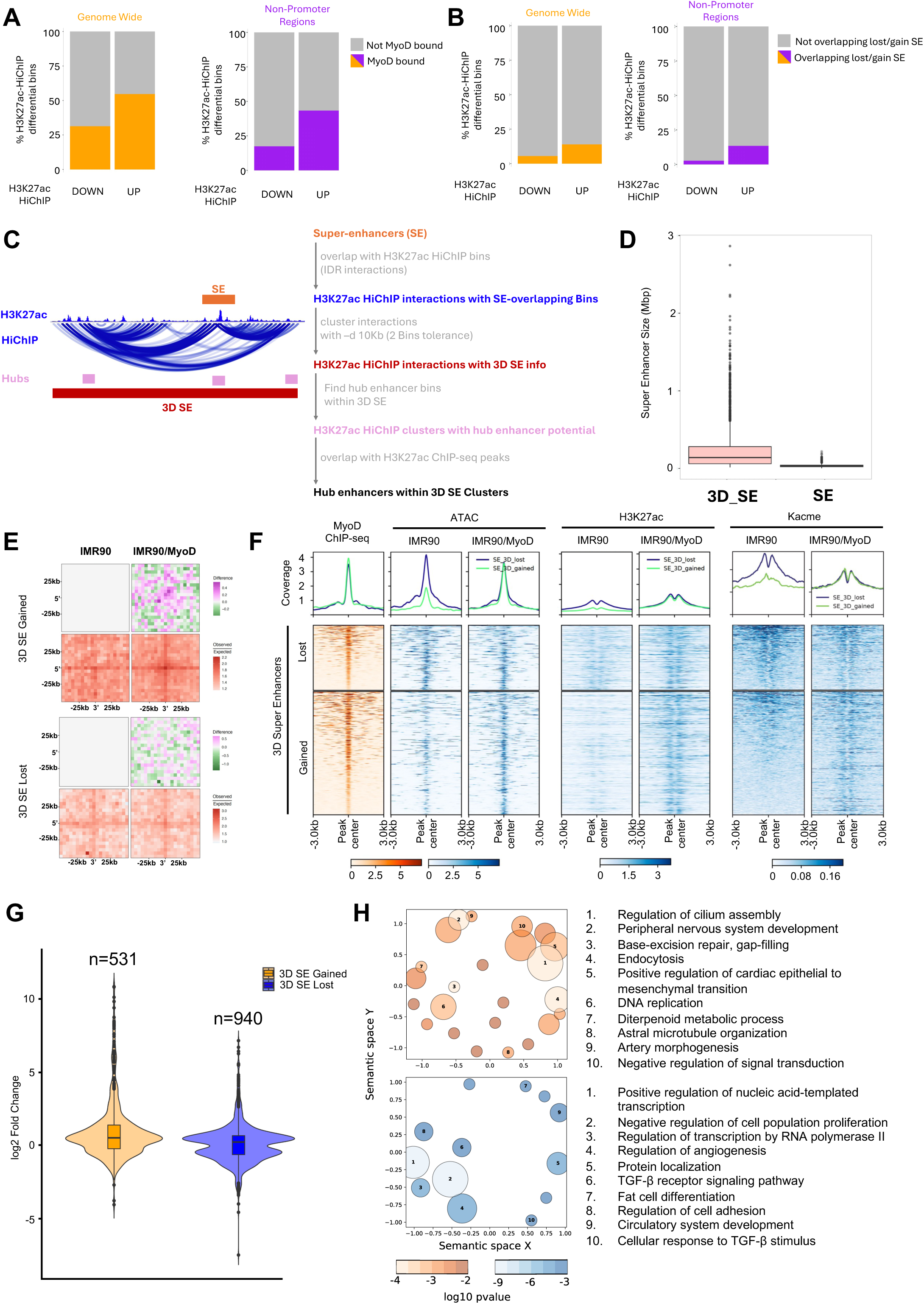
Identification of 3D SEs targeted by MYOD. a) Percentage of H3K27ac HiChIP differential bins bound or not by MyoD at the genome-wide level (orange) or specifically at non-promoter regions (purple). Differential HiChIP interactions were defined as p-value<0.05. b) Percentage of H3K27ac HiChIP differential bins overlapping or not SE at the genome-wide level (orange) or specifically at non-promoter regions (purple) bound by MyoD. c) Strategy to define 3D SEs. Hubs have been defined based on Huang et al., 2018. d) Box plot of size distribution of 3D SEs and linear SEs. e) Heatmap of Aggregate Peak Analysis (APA) of H3K27ac-HiChIP data centered at gained (top panels) or lost 3D SE (bottom panels) in IMR90 (left), IMR90/MyoD (right). f) Tornado and aggregate signal plots from left to right: MyoD ChIP-seq (orange) and ATAC-seq, H3K27ac ChIP-seq, and Kacme ChIP-seq (blue) in IMR90 and IMR90/MyoD. g) Box plots of log fold change of differentially expressed genes whose promoters interact with gained (orange) or lost (blue) 3D SEs through H3K27ac HiChIP loops. h) Gene Ontology of biological processes of up-and down-regulated DEGs in (g).

### MYOD binds and decommissions super-enhancers (SEs) of cell-of-origin and alternative lineage genes during human fibroblast trans-differentiation into skeletal muscle cells

As a large number of genome-wide MYOD chromatin binding events coincide with decreased chromatin accessibility also at genomic elements distal from promoters, we investigated the possibility that MYOD might also repress gene expression from non-promoter, distal genomic elements. Integration of MYOD ChIP-seq and ATAC-seq datasets showed that MYOD binding at non-promoter elements can result in either increased or decreased chromatin accessibility (Ext. Fig. 5A and B) that, again, were invariably associated to increased levels of H3K27 acetylation (Ext. Fig. 5C and D). Integration with CUT&RUN analysis of histone modifications further showed that MYOD-bound distal elements exhibiting increased chromatin accessibility were especially enriched in marks of enhancer identity (H3K4me1) and activation (H3K27ac) (Ext Fig. 4C). A slight increase in promoter identity (H3K4me3) was also observed at MYOD-bound distal elements with increased chromatin accessibility (Ext Fig. 4C). In contrast, reduced chromatin accessibility at MYOD-bound distal elements, again, did not coincide with any appreciable change in histone marks, except for a slight increase in H3K27ac (Ext Fig. 4D). An additional feature identified by this analysis was a trend of identity shift for MYOD-bound active enhancers into active promoters (Ext Fig. 4E). Interestingly, MYOD-bound non-promoter elements with increased chromatin accessibility coincided with formation of euchromatin and enhancer activation (Ext. Fig. 6A); instead, peaks with decreased chromatin accessibility exhibited a tendency toward loss of enhancer identity, despite retaining marks of euchromatin (Ext. Fig. 6A). As enhancers are the most relevant distal regulatory elements, and because we have observed a partial loss of enhancer identity in IMR90-MYOD cells (Ext. Fig. 4; Fig. 2E), we decided to focus on MYOD-bound enhancers, as previously reported^51^. In this regard, we were especially interested in the dynamics of super-enhancers (SEs) formation/decommissioning upon MYOD expression in IMR90 cells, as SE are typically implicated in the regulation of lineage identity genes^52^. We therefore first determined whether MYOD chromatin binding at non-promoter, distal genomic elements, coincided with SEs, including those already present in IMR90 cells and those formed in IMR90-MYOD cells. A total of 1674 SEs were identified in both IMR90 and IMR90-MYOD cells, using ROSE^52^. MYOD bound 78% (810 out of 1041) of SEs detected in IMR90 cells and 91% (1223 out of 1352) of SEs detected in IMR90-MYOD (Ext. Fig. 6B and C). While MYOD-bound SEs with increased chromatin accessibility were highly enriched with myogenic E-box motifs, MYOD-bound SEs with decreased chromatin accessibility were enriched with non-E-box motifs, mostly belonging to the Jun/Fos family members AP1 binding sites (Ext. Fig. 6D). Since SEs typically activate gene expression from a distance^53^, we used high-resolution (4KB) Hi-C datasets previously generated in IMR90, IMR90 GM, and DM conditions^28^ to identify SE loops with cognate promoter(s). Hi-C-based capture of SEs revealed the dynamics of enhancer-promoter (E/P) loops during the transition from IMR90 fibroblasts to IMR90-MYOD cells, discriminating newly formed vs lost SEs (Fig. 3A). Interestingly, this analysis also revealed several conserved SEs that were identified both in IMR90 and IMR90-MYOD cells (Fig. 3A). The dynamics of these conserved SEs was further investigated by Aggregate Region Analysis (ARA) of Hi-C data in IMR90 and IMR90-MyoD cultured in GM for 24 hours and then exposed to differentiation medium (DM) for additional 24 hours. This analysis illustrates how conserved SE between IMR90 and IMR90-MYOD cells are eventually lost along with IMR90-MYOD cell transition from culture in GM to terminal differentiation upon exposure to DM (Ext. Fig. 6E).

**Fig. 5.**
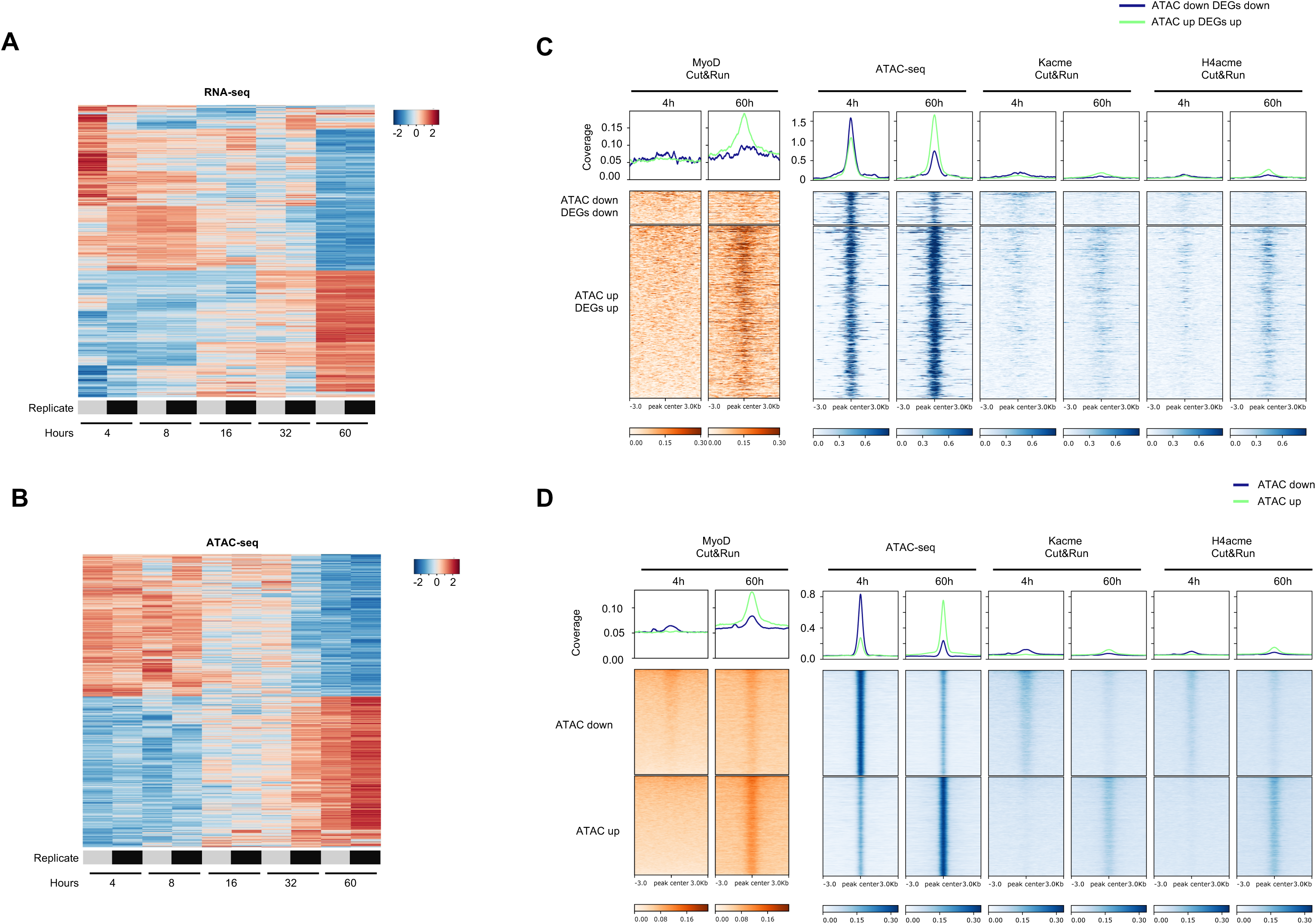
Opposite patterns of chromatin accessibility and Kacme levels at *MyoD*-bound loci during MuSCs activation. a-b) Heatmaps of differential gene expression (a) and promoter accessibility (b) of mouse satellite cells at 8, 16, 32 until 60 hrs post injury (all compared to 4 hrs; FDR-adjusted P < 0.05). c-d) Aggregate Plots (top panels) and Tornado plots (bottom panels) of CUT&RUN and ATAC-seq experiments in mouse satellite cells at 4 hrs (left) and 60 hrs (right) post injury overlapping promoter regions of DEGs (c) and non-promoter regions (d). From left to right: *MyoD*, ATAC-seq, Kacme, H4Kacme.

**Fig. 6.**
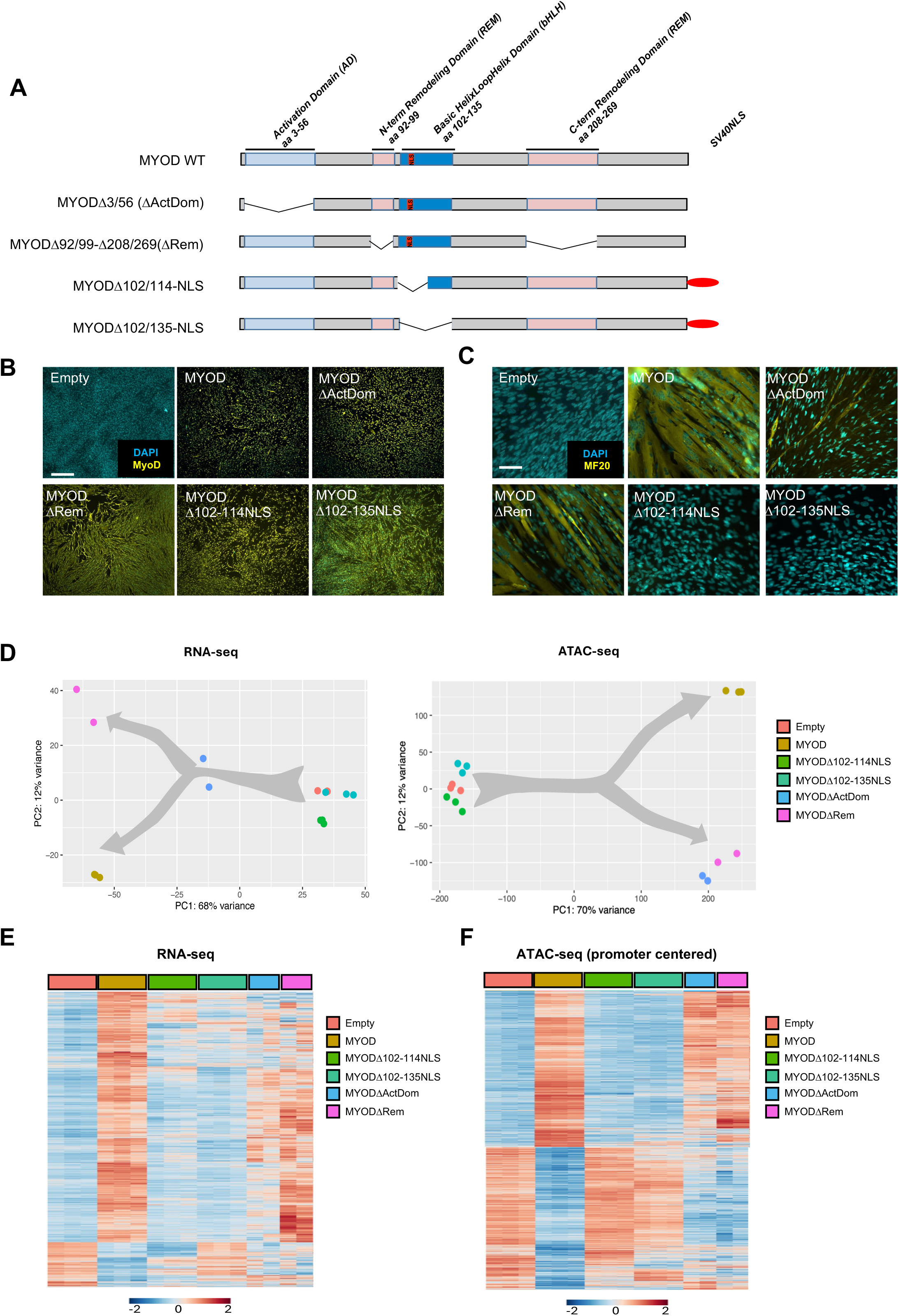
MYOD-mediated repression requires the integrity of functional domains previously implicated in MYOD-mediated activation of gene expression. a) Schematic representation of MYOD protein with indicated key functional domains and the MYOD deficient mutants used in the analysis. b) Representative image of immunofluorence analysis in for MyoD (yellow) in IMR90 cells ectopically expressing the different mutants described in (a) in growth condition. c) Representative image of immunofluorence analysis in for MF20 (yellow) in IMR90 cells ectopically expressing the different mutants described in (a) in differentiation condition. d) Principal Component Analysis (PCA) of gene expression data (top) and chromatin accessibility (bottom) MYOD mutants expressing lines in growth conditions. e-f) Heatmaps of differential gene expression (e) and promoter accessibility (f) of MYOD mutants expressing lines in growth conditions. FDR-adjusted P < 0.05 for RNA-seq and ATAC-seq, also |log2FC|>1.5 for ATAC-seq.

We next used MYOD ChIP-seq data to track the dynamics of MYOD-bound gained or lost SEs. MYOD-bound gained SE exhibited an increased Hi-C signal during the transition from IMR90 to IMR90-MYOD cells (Fig. 3B). In contrast, MYOD-bound lost SEs exhibited a drastic reduction in Hi-C signal related to chromatin interactions that define hubs of contacts typical of SEs, also referred to as frequently interacting regions (FIREs)^54^. Further integration with ATAC-seq data showed that MYOD-bound gained SEs exhibited increased chromatin accessibility (Fig. 3C), were typically marked by histone marks of active enhancers and euchromatin (Fig. 3D, top) and were enriched with myogenic E-box motifs (Fig. 3E, top). Conversely, MYOD-bound lost SEs exhibited a slight decrease in chromatin accessibility (Fig. 3C), were typically marked by histone marks of non-enhancer identity, although they retained marks of euchromatin (Fig. 3D, bottom), and were enriched with non-E-box motifs, mostly belonging to the Jun/Fos family members AP1 binding sites (Fig. 3E, bottom). Integration of Hi-C and RNA-seq data enabled the identification of DEGs downstream of promoters looping with MYOD-bound SEs. This analysis revealed distinct patterns of association between lost SEs with gene repression and reduced chromatin accessibility and gained SEs with gene activation and increased chromatin accessibility (Fig. 3F). Gained or lost SEs showed a trend of increased or decreased chromatin accessibility, respectively (Fig. 3G). Hi-C analysis of SE-associated promoters revealed that genes regulated by gained SE belong to biological processes related to general aspects of cellular differentiation and commitment to the myogenic lineage (Fig. 3H, top). In contrast, genes regulated by lost SEs belong to biological processes related to repression of growth factor-induced intracellular signaling (Fig. 3H, bottom).

Overall, these data define two types of MYOD-bound SEs. Gained SEs were not present in IMR90 cells and were generated upon MYOD expression via MYOD binding to E-box motifs at previously silent loci, according to a well-established sequence, by which MYOD targets compacted chromatin at nucleosomes in cooperation with pioneer factors, such as Pbx1/Meis, to promote chromatin remodeling and accessibility^5^. Lost SEs were present in IMR90 prior to the expression of MYOD and became decommissioned/inactivated upon MYOD binding to non-E-box motifs, via compaction and reduction of chromatin accessibility.

Because SEs consist of multiple hubs of chromatin interactions marked by H3K27ac, we performed HiChIP with H3K27ac antibodies and detected MYOD-bound HiChIP bins (Fig. 4A); however, only few of these MYOD-bound differential chromatin interactions marked by H3K27ac were found to overlap with SEs, as called by the traditional ROSE method (Fig. 4B). Thus, we sought to devise a novel approach to identify MYOD-bound 3D SEs, by using ROSE-derived SEs as starting point, and then overlapping linear SEs with the H3K27ac HiChIP bins, followed by clustering of all bins belonging to those interactions. Within the 3D SEs, we called hubs, according to Huang *et al.*, 2018^55^, thereby identifying bins within the 3D SEs that participate to more interactions, as compared to the genome-wide average (workflow illustrated in Fig. 4C). To further refine the hubs calls, we overlapped those bins with H3K27ac ChIP-seq peaks, which narrows the identification of 3D SEs regulatory hotspots (Fig. 4C). This method enabled the capture of considerably larger SEs, as compared to the conventional call of SEs used before (Fig. 4D) and revealed clear trends of increase (3D SE gained) or decrease (3D SE lost) from IMR90 to IMR90-MYOD cells (Fig. 4E). Importantly, while MYOD-bound gained SEs exhibited increased chromatin accessibility and H3K27ac signal, MYOD-bound lost SEs showed decreased chromatin accessibility, yet retained high levels of H3K27ac signal (Fig. 4F). The unexpected retention of high H3K27ac activation mark at MYOD-decommissioned SEs mirrors a similar phenomenon observed for MYOD-bound promoters of repressed genes shown in Fig. 1, thereby revealing the uncoupling of reduced chromatin accessibility and H3K27ac levels, as an unexpected, general feature of gene repression by MYOD from either promoters or enhancers. At the same time, this finding prompts the question of whether alternative histone modifications might provide a mark of MYOD-mediated gene repression. While most of the known histone marks did not show any pattern of association with MYOD-bound SEs that undergo chromatin compaction and direct repression of target genes, we turned our attention on a newly identified mark of histone H4 lysine 5 and 12 methylation and acetylation on the same side chain (H4K5/12 Kacme)^44^, as potential dynamic signal that could associate with MYOD-mediated inactivation of SEs. Indeed, ChIP-seq with Kacme antibodies showed a clear enrichment at MYOD-bound SEs with increased chromatin accessibility, while a reduction in Kacme signal marked MYOD-bound SEs with decreased chromatin accessibility (Fig. 4F). Upon integration of 3D SEs with RNA-seq data, by HiChIP H3K27ac-detected loops, we identified a large amount of DE genes downstream to the promoters looping with these SEs (Fig. 4G). Gene ontology analysis revealed that SE-upregulated genes were mostly related to biological processes of cellular commitment and differentiation toward skeletal muscle lineage, while SE-downregulated genes referred to processes related to repression of alternative cell lineages, in addition to genes implicated in cell proliferation (Fig. 4H). Examples of genes downregulated upon MYOD binding and decommissioning of SEs are shown in Ext. Fig. 7A and B.

Another feature of SEs is the frequent enrichment in CTCF-marked chromatin interactions^54^. As previous studies have revealed an association between MYOD and CTCF chromatin binding^29^, we also performed HiChIP with CTCF antibodies to investigate whether MYOD binding alters CTCF-mediated chromatin interactions at SEs. This analysis identified a coherent trend of reduction in the strength of CTCF-mediated chromatin interactions at lost SEs, while an opposite pattern was observed at gained SEs (Ext. Fig. 7C). Integration with MYOD and Kacme ChIP-seq and ATAC-seq revealed that MYOD-bound and lost SEs with reduced strength of CTCF-mediated chromatin interactions exhibited reduced chromatin accessibility and Kacme signal. On the contrary, the opposite pattern was observed with MYOD-bound, gained SEs with increased strength of CTCF-mediated chromatin interactions (Ext. Fig. 7D). Surprisingly, in both cases MYOD binding to SEs led to increased CTCF chromatin binding, indicating that CTCF chromatin affinity/binding can be dissociated from CTCF-mediated strength of chromatin interactions.

Overall, the data support the identification of a novel property of MYOD, as repressor of gene expression during somatic cell trans-differentiation into skeletal muscle, via direct binding to promoters and distal elements (enhancers and SEs), through a common mechanism that entails binding to non-E-box elements, invariably leading to reduction of chromatin accessibility, with decrease in strength of CTCF-mediated chromatin interactions and Kacme levels at SEs.

### Opposite patterns of chromatin accessibility and Kacme levels at MyoD-bound loci during muscle stem cell (MuSC) activation

We next investigated whether MYOD-mediated gene repression could also be observed in physiological conditions. To this purpose we turned our attention to two specific stages of skeletal muscle regeneration in mice – namely 4-and 60-hours post-injury. These timepoints were selected among sequential time-points across MuSCs activation, since they coincided with lack or induction of *MyoD* expression, respectively, as reported by Dong et al.^34^. RNA-seq and ATAC-seq analysis revealed coherent and parallel patterns of both gene upregulation with increased chromatin accessibility at their promoters and gene downregulation with reduced chromatin accessibility at their promoters (Fig. 5A and B). CUT&RUN analysis of genome-wide chromatin binding of *MyoD* and enrichment in Kacme showed that at 4 hours no MyoD binding was detected at promoters of DEGs with differences in chromatin accessibility (Fig. 5C and D), consistent with the lack or very low levels of *MyoD* expression at this stage^34^. However, we detected *MyoD* recruitment at these promoters in MuSCs isolated at 60 hours post-injury, accompanied by a consensual reduction in Kacme levels at promoters of repressed genes with reduced chromatin accessibility, and increased Kacme levels at promoters of activated genes with increased chromatin accessibility, using antibodies that recognize Kacme in any context (pan-Kacme), or ones that are specific for Kacme on histone H4 on K5/12 (H4Kacme) (Fig. 5C). Likewise, *MyoD* enrichment at distal elements showing reduced chromatin accessibility was associated with reduced Kacme levels, whereas *MyoD*-bound distal elements with increased chromatin accessibility showed enrichment in Kacme levels (Fig. 5D). These data further support the conclusion that MYOD either activates or represses gene expression in muscle progenitors, by increasing or reducing chromatin accessibility and Kacme levels, respectively, within the context of muscle regeneration. Tracks of representative loci are shown in Extended Figure 8.

### MYOD-mediated repression requires the integrity of functional domains previously implicated in MYOD-mediated activation of gene expression

Our data reveal a “logic” connection linking E-box independent MYOD chromatin and gene repression from promoters of mitogen/growth factor-responsive genes implicated in the activation of the cell cycle and from SEs of cell-of-origin and alternative cell lineage genes. Of note, the mechanism of MYOD-mediated gene repression implicates events that are opposite to those implicated in MYOD-mediated gene activation, raising the issue of whether distinct functional domains of MYOD are involved in these different tasks. For this reason, we used MYOD mutants in which specific functional domains are deleted. Figure 6 illustrates these MYOD mutants, which include deletion (Δ3-56) of the N-terminal activation domain (ΔActDom); small (Δ102-114) and large (Δ102-135) deletion of the first helix (note that since this deleted fragment contains the nuclear localization signal (NLS), a heterologous NLS was added to induce nuclear localization of these mutants); deletion of both N terminal (Δ92-98) and C-terminal (Δ208-269) chromatin remodeling domains (ΔRem) (Fig. 6A). These mutants were expressed in IMR90 cells in a doxy-regulated manner, as for the MYOD WT, and they all invariably showed a nuclear localization upon culture in GM, without any noticeable changes in cell morphology and induction of the endogenous MYOD transcript (Fig. 6B and Ext. Fig. 9A); however, when incubated in DM, while both IMR90-MYODΔ102-114NLS and IMR90MYODΔ102-135NLS cells completely fail to differentiate into multinucleated, MyHC-positive myotubes, and IMR90-MYODΔActDom cells formed very sporadic myotubes, IMR90-MYODΔRem showed an impaired differentiation index, as compared to IMR90-MYOD WT, yet eventually formed multinucleated MyHC-positive myotubes, albeit with lower efficiency and reduced caliber (Fig. 6C). Thus, in principle, this evidence indicates that mutations that disrupt domains required for activation of gene expression, such as chromatin remodeling, might ultimately be tolerated/compensated to allow the formation of terminally differentiated muscles.

We performed parallel RNA-seq and ATAC-seq in the above-mentioned cells cultured in GM for 24 hours after doxy-induced expression of MYOD WT or mutants. This analysis showed mutant-specific patterns of gene expression and chromatin accessibility at promoters of target genes, whereby IMR90-MYODΔ102-114NLS and IMR90-MYODΔ102-135NLS exhibited profiles very similar to IMR90 control cells, while IMR90-MYODΔActDom and IMR90-MYODΔRem cells showed “intermediate” profiles between IMR90-MYOD WT and IMR90 control cells (Fig. 6D-F). While these “intermediate” profiles of gene expression and chromatin accessibility are mostly accounted by the partial ability of IMR90-MYODΔActDom and IMR90-MYODΔRem to activate gene expression and chromatin remodeling, a closer inspection revealed individual and common patterns of gene repression and reduction in chromatin accessibility among the mutants that define distinct modules of MYOD-mediated repression (Ext. Fig. 8A). We identified one cluster of genes, whose repression requires the 114-135aa sequence, as they are repressed by all mutants except MYODΔ102-135NLS (Fig. 6E – and Ext. Fig. 9B). Gene ontology analysis revealed that these genes are all responsive to mitogens and implicated in the activation of the cell cycle (Ext. Fig. 9C). One cluster of genes, whose repression required only the 102-114aa sequence, included growth factor-responsive genes (Fig. 6E and Ext. Fig. 9D). Another subset of MYOD-repressed genes required the entire 102-135aa sequence, as they are not repressed by either MYODΔ102-114NLS or MYODΔ102-135NLS, but continued to be repressed by MYODΔActDom or MYODΔRem, and included growth factor-responsive genes also implicated in cell adhesion and fusion (Fig. 6E and Ext. Fig. 9C and D). A subset of MYOD-repressed genes required the AD and Rem domains, as they are not repressed by either MYODΔActDom or MYODΔRem, but continued to be repressed by MYODΔ102-114NLS or MYODΔ102-114NLS. These genes were implicated in cell-of-origin or alternative cell lineages (Fig. 6E and Ext. Fig. 9F-H).

## Discussion

The results shown here reveal a novel property of MYOD as transcriptional repressor from non-E-box sequences, thereby challenging the existing dogma that has historically restricted MYOD biological property to that of a sequence-specific transcriptional activator from myogenic E-box sequences. As such, this finding extends, in principle, our knowledge on MYOD from a monotone activator of gene expression to versatile regulator of gene expression.

We show that MYOD exerts its function as a dual activator and repressor of gene expression during the process of cellular trans-differentiation into skeletal muscle, by coordinating two key events within the nuclear reprogramming toward the myogenic lineage - activation of the genes that establish the new cell identity (*i.e.*, skeletal muscle lineage), and erasure of the cell-of-origin lineage genes. This function is supported by previous works reporting on alternative mesenchymal lineages adopted by MYOD-deficient MuSCs^56–58^. Likewise, extra-ocular muscles, in which repression of mesodermal genes is incomplete, show a reduced expression of MYOD^59^. Because nuclear reprogramming is the fundamental process for determination of cell lineage identity during development and adult tissue regeneration, as well as for induced pluripotency and for somatic cell trans-differentiation^60^, our results might provide a general paradigm for a dual function extended to other tissue-specific transcription factors (TFs) implicated in nuclear reprogramming.

MYOD-mediated repression of transcription also includes the downregulation of growth factor-, cytokine-, and mitogen-responsive genes, suggesting that MYOD might prevent a promiscuous activation of gene expression in muscle progenitors exposed to a multitude of extracellular signals. This condition typically occurs when muscle progenitors are exposed to developmental or regeneration signals during embryonal and adult skeletal myogenesis, respectively. In particular, during skeletal muscle regeneration, the activation of MuSCs entails loss of anatomical insulation from the surrounding environment that is provided by the myofiber basal lamina (otherwise, defined as quiescence niche). Consequently, activated MuSCs are exposed to a plethora of regeneration cues and signals, which would promiscuously activate gene expression if not properly interpreted. In this regard, we propose that MYOD functions as a “storm shelter” to prevent the expression of regeneration-activated genes that could eventually bias MuSC function. This is particularly important when activated MuSCs are challenged by the inflammatory cytokines released during the initial stages of regeneration, by M1 macrophages and other immune cells, which would otherwise interfere with MuSC commitment and differentiation into multinucleated myofibers^61^. Furthermore, the transition of MuSCs from proliferation to cell cycle withdrawal prior to their differentiation into myofibers also requires coordinated expression of genes that regulate different phases of cell cycle, within the complexity of the regeneration milieu. We propose that MYOD-mediated repression of gene expression provides transcriptional tolerance and competence for proper response to regeneration cues, to coordinate sequential patterns of gene expression in MuSCs along their transition from activation to differentiation into myofibers.

Of note, we identified specific molecular modules used by MYOD to repress gene expression from promoters or enhancers of target genes with dedicated domains, whereby repression of mitogen-and growth factor-responsive genes from promoters requires a highly conserved domain within the first helix, while repression of cell-of-origin and alternative lineage genes from SE requires the AD and CRDs.

Our results reveal an interplay between genetic, epigenetic, and molecular determinants that confers on MYOD a dual function of transcriptional activator or repressor. The genetic determinant that discriminates between these two functions is the alternative MYOD chromatin recruitment through binding to myogenic E-box or non-E-box motifs, which leads to two opposite patterns of chromatin accessibility and histone modifications. Previous works established that MYOD-mediated activation of gene expression from E-box motifs entails a sequence of events prompted by the reported ability of MYOD to bind nucleosomes at previously silent loci in cooperation with pioneer factors, such as Pbx1/Meis^5,26^, followed by increased chromatin accessibility, which allows full recognition and binding to myogenic E-box sequences^13–16^. Heterodimerization with E2A gene products, E12 and E47, enables MYOD to fully activate target gene expression^19^. Specific domains confer on MYOD the property as an E-box-specific transcriptional activator – namely, the basic domain that restricts its DNA binding affinity to specific E-box motifs, the HLH domain that promotes interactions with E12/47an acidic activation domain (ActDom) at the N-terminus, and two chromatin remodeling domains located at the C/H-rich domain and C-terminus^21^. Previous studies established that binding to E-box sequences triggers intramolecular changes in MYOD that unlock the ActDom to adopt a conformation to activate transcription^23,62^. Likewise, binding to nucleosomes promotes the chromatin remodeling activity of bHLH TFs and activates the enzymatic activity of transcriptional co-activators, such as acetyltransferases^63,64^. Thus, the evidence that MYOD chromatin recruitment at non-E-box motifs within accessible chromatin promotes the opposite outcome – chromatin compaction and repression of gene transcription – reveals a hitherto unappreciated functional versatility of MYOD that is imparted by the alternative chromatin recruitment through genetic determinants - E-box or non-E-box motifs. This evidence also suggests that chromatin recruitment via E-box-independent interactions might turn functional domains of MYOD into effectors of MYOD-mediated repression.

MYOD-mediated repression of gene expression occurs mostly when muscle progenitors are exposed to growth factors, cytokines, and mitogens, as well illustrated by the model of IMR90/MYOD cells cultured in GM. This condition is incompatible with MYOD heterodimerization with E2A proteins and productive binding to E-box motifs^65,66^. Likewise, MYOD is expressed in MuSCs few hours after their activation – a stage that coincides with MuSC proliferation, which is incompatible with the activation of muscle gene expression - thus suggesting that MYOD might exert functions alternative to the activation of muscle gene expression. We propose that at this stage MYOD pervasively binds the genome through weak interactions at both E-box and non-E-box motifs. Initial binding to E-box motifs primes promoters and enhancers of muscle genes for subsequent activation of gene expression, upon exposure to pro-differentiation cues, as proposed by previous works^67^, and supported by recent evidence^68^. Within this context, transient recruitment of transcriptional co-repressors holds E-box-driven activation of muscle genes until the exposure to pro-differentiation cues, as a mechanism that warrants a timely and coordinated activation of muscle-gene expression during myoblast to myotube transition^37–40,67^. In this regard, our discovery that MYOD represses the expression of cell-of-origin and alternative lineages or growth factor-inducible genes from non-E-box motifs substantially differs from the transient inhibition of transcription from E-box sequences. We have identified reduced chromatin remodeling, decreased levels of Kacme and strength of CTCF-mediated chromatin interactions as key epigenetic determinants of MYOD-mediated gene repression from non-E-box motifs. However, we did not detect any marker of constitutive heterochromatin at MYOD-bound non-E-box within promoters/enhancers of repressed genes. It is possible that formation of constitutive heterochromatin at these loci occurs later during the process of muscle differentiation, possibly instigated by MYOD-mediated activation of the additional transcriptional repressors, such as the zinc-finger protein RP58 (also known as Zfp238)^41^ or DNMT3^69^.

In sum, our data extend the biological properties of MYOD beyond our current knowledge, by revealing its dual function as activator and repressor of gene expression, through a multifaceted mechanism. The selection of either one of these functions is determined by an interplay between genetic, epigenetic, and molecular determinants upon muscle progenitor transition through cell states and exposure to regeneration cues. This is consistent with the early prediction that MYOD senses and integrates many facets of cell state^70^. We argue that defective execution of MYOD-mediated repression of gene expression might be tolerated, as long as potential mutations that impair this property do not affect MYOD-ability to activate skeletal myogenesis. We postulate that one consequence of this trade-off could be the defective long-term maintenance of the functional properties of myofibers, as they represent a perennial tissue with limited nuclear turnover that relies on MuSC-mediated regeneration.

## Methods

### Cell Culture Experiments

IMR90-EMPTY or MYOD WT or mutant-expressing IMR90- cells were maintained in growth media (GM) consisting of EMEM (ATCC) supplemented with 10% FBS (Omega Scientific). Cells were regularly tested for absence of mycoplasma.

Myogenic conversion. Myogenic conversion was performed as previously described in Dall’Agnese et al 2019^28^.

### Antibodies

The following commercially available primary antibodies were used in this study: mouse monoclonal anti-MYOD (BD Bioscience, Cat #554130), recombinant abflex anti-H3K27ac (Active Motif, Cat #91193), mouse monoclonal anti-MyHC (DSHB, MF-20), goat polyclonal anti-GATA6 (BioTechne, Cat #AF1700), anti-CTCF (Active Motif Cat #91285), anti-CTCF (CST Cat#3418S), anti-H3K4me3 (CST Cat #9751), anti-H3K4me1 (Active Motif Cat #39635), anti-H3K27me3 (Active Motif Cat #39155), anti-H3K9me3 (Active Motif Cat #39161), H4Kacme and Kacme antibodies were previously described^44^. The secondary antibodies were goat anti-mouse IgG, Fc subclass 1 specific Cy3-conjugated (Jackson ImmunoResearch, 115-545-207), goat anti-mouse IgG, Fc subclass 2b specific 488-conjugated (Jackson ImmunoResearch, 115-165-205), donkey anti-mouse IgG 568 (Life Technologies, ref A10037) and donkey anti-goat 488 (Life Technologies, ref A32814).

### Immunofluorescence

Cells were fixed with 4% PFA in PBS, permeabilized with 0.5% TX100 and blocked with 5% BSA in PBS. Cells were stained with anti-MYOD (BD Bioscience, Cat #554130) and anti-myosin heavy chain (DSHB, MF20) or anti-MYOD (BD Bioscience, Cat #554130) and anti-GATA6 (BioTechne, AF1700) O/N at 4C followed by anti-mouse IgG, Fc-subclass 2b 488 conjugate (Jackson ImmunoResearch) and anti-mouse IgG, Fc-subclass 1 Cy3 conjugated (Jackson ImmunoResearch) or Donkey anti-mouse Alexa 568 (Life Technologies, ref A10037) and Donkey anti-goat Alexa 488 (Life Technologies, ref A32814) respectively for 1 hr at RT in the dark. Nuclei were then counterstained with 2 ug/ml Hoechst 33258 pentahydrate (bis-benzimide) (Life Technologies) 5 minutes at RT. Images were acquired with fluorescence microscope. Fields reported in figures are representative of all examined fields.

### Mouse injury and muscle stem cells isolation

Muscle injury and MuSCs FACS isolation was performed as previously described in Dong et al 2022^34^. All animal experiments were approved by the HKUST Animal Ethics Committee.

### mRNA expression analysis

Total RNA was extracted using Quick-RNA Microprep KIT (Zymo Research, R1051) following manufacture’s recommendation. RNA concentration was measured on Qubit (Invitrogen). 100-500 ng of RNA was reverse transcribed using High-Capacity cDNA Reverse Transcription Kit (Applied Biosystems, 4368813). Real-time quantitative PCR (qPCR) was performed using Power SYBR Green Master Mix (Life Technologies) following manufacture’s indications. Expression was normalized to *TBP* for IMR90 cells using 2^-ΔΔCt^ method. Primers used in the study: hMYOD 5’-TTAACCACAAATCAGGCCGG-3’ 5’-CAAAGTGCTGGCAGTCTGAATG-3’, mMYOD 5’-AGCACTACAGTGGCGACTCA-3’ 5’-GGCCGCTGTAATCCATCAT-3’, GATA6 5’-AGAAGCGCGTGCCTTCATC-3’ 5’-TTTCTGCGCATAAGGTGGT-3’, TNNT2 5’-TCAAAGTCCACTCTCTCTCCATC-3’ 5’-GGAGGAGTCCAAACCAAAGCC-3’, ITGA7 5’-TCGAACTGCTCTTCTCACGG-3’, 5’-CCACCAGCAGCCAGCTC-3’, FN1 5’-CTGGAACCGGGAACCGAATA-3’ 5’-CGAAAGGGGTCTTTTGAACTGT-3’, MYH3 5’-CGAAGCTGGAGCTACTGTAA-3’ 5’-CCATGTCCTCGATCTTGTCATA-3’, IL6 5’-CGGGAACGAAAGAGAAGCTCTA-3’ 5’-GGCGCTTGTGGAGAAGGAG-3’, cFOS 5’-CAGACTACGAGGCGTCATCC-3’ 5’-TCTGCGGGTGAGTGGTAGTA-3’ TBP 5’-GCGCAAGGGTTTCTGGTTTG-3’ 5’-GTAAGGTGGCAGGCTGTTGT-3’.

### HiChIP H3K27ac and CTCF library preparation

HiChIP experiments were performed using the ARIMA HiC+ kit according to manufacturer’s protocol. Briefly, 5 million IMR90 or IMR90/MyoD were resuspended in 5ml of growth media and fixed with 2%FA for 10 minutes at room temperature followed by quenching with STOP solution1 according to ARIMA HiC+ protocol. Digestion was performed overnight at 37C. Biotinylated enriched fragments were pulldown with anti-H3K27ac (Active Motif Cat #91193) or anti-CTCF (Active Motif Cat #91285). H3K27ac or CTCF enriched material was then used as input for library preparation using Accel-NGS 2S Plus DNA Library Kit (Swift Biosciences Cat #21096) according to ARIMA HiChIP library preparation protocol. Obtained libraries were sequenced on NOVASeqS4 PE 2×100 at the IGM UCSD at a depth of ∼300M per library.

### CTCF ChIP-seq library preparation

For each ChIP-seq replicate, 4×10^6^ cells with 1% FA for 10 minutes at RT followed by quenching with glycine (final concentration 200 mM) for 15 minutes on ice. Nuclei were then extracted in hypotonic buffer (10 mM Tris-HCl, pH 8.0, 10 mM NaCl, 0.05% NP-40, 1 mM PMSF and 1x protease inhibitor) and lysed in lysis buffer (50 mM Tris-HCl, pH 8.0, 150 mM NaCl, 5 mM EDTA, pH 8.0, 0.5% SDS, 0.5% NP-40, 1 mM PMSF and 1x protease inhibitor). Chromatin was sheared with sonicator (S2 Covaris) to an average DNA fragment length of 200-500bp. 15µg of sonicated chromatin were diluted in 500µl RIPA Dilution Buffer (50 mM Tris-HCl, pH 8.0, 150 mM NaCl, 1 mM EDTA, pH 8.0, 0.1% Sodium Deoxycholate, 0.7% NP-40, 1 mM PMSF and 1x protease inhibitor), and precleared for 3 hours at 4C with Protein A/G dinabeads. In parallel, 10µl of CTCF antibody (CST Cat#3418S), or 1µg of rabbit IgG (Santa Cruz Cat #sc-2027) were prebound to Protein A/G dinabeads in 500µl of PBS containing 5 µg/ml BSA, for 3 hours at 4C. Beads bound antibody were mixed with precleared chromatin and incubated O/N at 4C on a rotator. Chromatin bound fraction was washed with RIPA washing buffer (50 mM Tris-HCl, pH 8.0, 150 mM NaCl, 1 mM EDTA, pH 8.0, 0.5% Sodium Deoxycholate, 1% NP-40) for 4 times, followed by LiCl washing buffer (10 mM Tris-HCl, pH 8.0, 250 mM LiCl, 1 mM EDTA, pH 8.0, 1% Sodium Deoxycholate, 1% NP-40), and TE buffer (10 mM Tris-HCl, pH 8.0, 1 mM EDTA), each wash for 10 minutes at 4C. Chromatin was then eluted in elution buffer (1% SDS in TE buffer) at 65C for 6 hours 600 RPM rotation. Precipitated material and 10ng of input were used as input for library preparation using Accel-NGS 2S Plus DNA Library Kit (Swift Biosciences Cat #21096) according to manufacturer’s protocol. Obtained libraries were sequenced on NOVASeqS4 PE 2×100 at the IGM UCSD at a depth of ∼100M per library.

### CUT&RUN library preparation

For each CUT&RUN experiment 100K IMR90 or 250K muscle stem cells were used as input. CUT&RUN experiments were performed using the CUT&RUN Assay Kit (CST Cat #86652) with the following modification: cells were briefly fixed with 0.1% FA for 2 minutes at RT followed by quenching with glycine; antibody binding was performed O/N at 4C. Antibodies used: anti-MYOD (BD Bioscience, Cat #554130), anti-H4Kacme, anti-Kacme, anti-H3K27ac (Active Motif Cat #91193), anti-H3K4me3 (CST Cat #9751), anti-H3K4me1 (Active Motif Cat #39635), anti-H3K27me3 (Active Motif Cat #39155), anti-H3K9me3 (Active Motif Cat #39161). CUT&RUN libraries have been generated using the NEBNext^®^ Ultra™ II DNA Library Prep Kit for Illumina (NEB Cat #E7645S) according to manufacturer’s protocol. Libraries were sequenced at a depth of ∼20M per library on NOVASeqS4 PE 2×100 at the IGM UCSD.

### ATAC-seq library preparation

100K freshly collected IMR90 cells were subjected to ATAC-seq library preparation using the ATAC-seq Kit (Active Motif Cat #53150) according to manufacturer’s protocol. Libraries were sequenced at a depth of ∼200M per library on NOVASeqS4 PE 2×100 at the IGM UCSD.

### RNA-seq library preparation

Equal inputs of total RNA (10-100 ng) were used to generate stranded total RNA libraries for sequencing using the Illumina® Stranded Total RNA Prep, Ligation with Ribo-Zero Plus (Illumina Cat #20040525). ERCC RNA Spike-in mix was added at the start of the protocol according to manufacturer’s instruction (Thermo Fisher Scientific Cat # 4456740). Libraries were sequenced at a depth of ∼50M per library on NOVASeqS4 PE 2×100 at the IGM UCSD.

### RNA-seq data analysis

Data were checked for quality with FASTQC (v0.11.9, available online at: http://www.bioinformatics.babraham.ac.uk/projects/fastqc/), reads were trimmed with Trimmomatic (v0.39)^71^ to eliminate low quality bases and adapters (parameters: PE - phred33 ILLUMINACLIP:NexteraPE-PE.fa:2:30:10:2:true MAXINFO:40:0.1 MINLEN:45), and aligned to the hg19 Ensembl (November 2015) version of the human genome with STAR (v 2.7.3a)^72^, using a custom genome and index to simultaneously map and quantify ERCC92 spike-in-derived reads (parameters: --runMode alignReads --runThreadN 12 -- readFilesCommand zcat --outSAMtype BAM SortedByCoordinate --quantMode GeneCounts). Counts data (from STAR output in column 4, based on library preparation strandedness: second-strand) from all conditions were filtered based on their raw count, keeping genes where the sum of the counts for all samples was higher than 10. Size factors were calculated using the R environment (v4.1.3) package DESeq2 (v1.34.0)^73^ estimateSizeFactors function - using the counts related to the spike-in transcripts as control genes, then normalised and logged with the rlog function. DESeq2 was also used to perform Principal Component Analysis (PCA) and differential gene expression analysis (significance threshold: BH FDR<0.05). Differential expression analysis results were visualized with the heatplot function (parameters: method=“complete”, labRow= FALSE, dend=“row”, returnSampleTree=F, zlim=c(-5,5)) from the made4 (v1.68.0)^74^ package, after expression values from the DESeq2 object were turned into a matrix of Z-scores. Gene Ontology was performed on down-and up-regulated genes separately, using the gene symbols as input in EnrichR (available online at https://maayanlab.cloud/Enrichr/)^75^, and gene sets repositories GO Biological Processes 2023 and Descartes Cell Types and Tissue 2021. Top10 gene ontology terms shown in figure panels were ordered by pvalue. Gene Ontology bubble plots to summarize top50 gene ontology terms, were generated with the Python (v.3.8.1) GO-Figure^76^ script (parameters:-j standard-c log10-pval-e 100-si 0.5-s members-g single-n bpo--font_size small-q svg).

### ATAC-seq data analysis

Data were checked for quality with FASTQC, reads were trimmed with Trimmomatic to eliminate low quality bases and adapters (parameters: PE-phred33 ILLUMINACLIP:NexteraPE-PE.fa:2:30:10:2:true MAXINFO:40:0.1 MINLEN:45), aligned to the hg19 UCSC (January 2016) version of the human genome with Bowtie2 (v2.3.5.1)^77^ (parameters:--no-unal--local--very-sensitive-local--no-discordant--no-mixed--dovetail--phred33), sorted and converted into BAM format with the SAMtools sort function. Peak calling was performed with the Macs2 (v2.2.6) ^78^ callpeak function (parameters:-f BAMPE-g hs-B-q 0.05--nomodel--shift-100--extsize 200) and filtered for black-list regions with the BEDtools suite (v2.29.2)^79^ intersect (parameters:-v), and sorted with the sort function (parameters: - k8,8nr). Irreproducible Discovery Rate (IDR) was calculated with the idr function (v2.0.3)^80^ (parameters: --input-file-type narrowPeak --plot --only-merge-peaks) to retain a single peaks list from each condition. Differential chromatin accessibility analysis was carried out in R, with the DESeq2 package. The R package ChIPQC (v1.30.0)^81^, (GetGRanges function was used to import peak lists from each biological replicate, a consensus peak list was calculated for all conditions using the IRanges (v2.28.0)^82^,function reduce, keeping peaks that were called in at least two samples. Counts for each peak in each biological replicate were quantified with the Rsubread (v2.8.2)^83^ featureCounts function (parameters: isPairedEnd= T, countMultiMappingReads= F, maxFragLength= 100). Counts were normalised and logged with the DESeq2 function rlog; DESeq2 was also used to perform Principal Component Analysis (PCA), and differential chromatin accessibility analysis (significance threshold: BH FDR<0.05). Differentially Accessible Regions (DARs) at gene promoter regions were retrieved with the IRanges function promoters (parameters: TxDb.Hsapiens.UCSC.hg19.knownGene, 1000, 200; mouse genome version was: TxDb.Mmusculus.UCSC.mm10.knownGene). Promoter DARs were visualized with the made4 function heatplot (parameters: method=“complete”, labRow= FALSE, dend=“row”, returnSampleTree=F, zlim=c(-5,5)), after expression values from the DESeq2 object were turned into a matrix of Z-scores. Due to R visualization limits, only promoters with |log2FC|>1.5 were plotted. Motif analysis was carried out with the HOMER (v4.11)^84^, findMotifsGenome function (parameters: hg19-size given-nomotif).

### CUT&RUN data analysis

Data were checked for quality with FASTQC, reads were trimmed with Trimmomatic to eliminate low quality bases and adapters (parameters: PE-phred33 ILLUMINACLIP:NexteraPE-PE.fa:2:30:10:2:true MAXINFO:40:0.1 MINLEN:45), aligned either to the hg19 UCSC (January 2016) version of the human genome (for the CUT&RUN for histone marks in IMR90 and IMR90/MyoD), or to the mm10 Ensembl (November 2019) version of the mouse genome (for the CUT&RUN of Kacme and *MyoD* in MuSCs), with Bowtie2 (parameters:--phred33--end-to-end--no-unal--local--very-sensitive-local--no-mixed--no-discordant-I 10-X 700 –dovetail) and converted to BAM format with the SAMtools (v1.10)^85^ function view (parameters:-bS). Peak calling was performed with the Macs2 callpeak function (parameters:-g hs--nomodel--SPMR--call-summits-p 0.05) using IgG chromatin as control, and filtered for black-list regions with BEDtools intersect (parameters:-v), and sorted with the Unix sort function (parameters:-k8,8nr). Irreproducible Discovery Rate (IDR) was calculated with the idr function (parameters:--input-file-type narrowPeak--plot--only-merge-peaks) to retain a single peaks list from each condition. ChromHMM (v1.24)^86^ BinarizeBed function (parameters:-b 200-peaks) was used to bin the genome in 200bp windows according to each IDR peak list. The BinarizeBed output was used as input to ChromHMM LearnModel (parameters:-b 200 12 hg19), to build a 12 chromatin states model based on the histone marks IDR peaks. Chromatin state numbers were converted to chromatin states names in R, based on the combination of histone marks they were enriched for.

### Hi-C, H3K27ac and CTCF HiChIP data analysis

Hi-C data analysis was previously described in Dall’Agnese et al 2019^28^. HiChIP data were checked for quality with FASTQC, and alignment, filtering, normalization, and loop calling were performed with the Arima version of MAPS (v2.0)^87^ bash script (parameters:-C 0-F 1-M 1-H 1-o hg19-f 1-s 5000-r 2000000-d 2-Q 30-l 1000). Irreproducible Discovery Rate (IDR) was calculated with IDR2D (available online at https://idr2d.mit.edu)^88^ parameters: value transformation= log additive inverse (-log(x)), ambiguity resolution method= overlap, remove non-standard chromosome, max gap= 1000, max factor= 1.5, jitter factor= 0.0001, mu= 0.1, sigma=1, rho=0.2, p=0.5, epsilon=0.001). Differential HiChIP analysis was performed with the DiffAnalysisHiChIP.r script within FitHiChIP (v7.1)^89^, and significance threshold was set to p-value<0.05.

### ChIP-seq data analysis

MyoD and H3K27ac ChIP-seq data analysis details are reported in Dall’Agnese et al., 2019^28^. CTCF ChIP-seq data were checked for quality with FASTQC and aligned to the hg19 UCSC (January 2016) version of the human genome with Bowtie2 (parameters:--very-sensitive-local). Peak calling was performed with the Macs2 callpeak function (parameters:-g hs-B--nomodel--SPMR-q 0.05), using input chromatin as control.

### Genome-wide tracks generation and visualization

Genome-wide tracks of RNA-seq signal in BEDGRAPH format (forward and reverse) were generated with the DeepTools (v3.5.4)^90^, function bamCoverage (parameters:--normalizeUsing CPM--effectiveGenomeSize 2864785220--binSize 100--filterRNAstrand forward or reverse--outFileFormat bedgraph--scaleFactor $sizeFactor) using as scale factors the size factors calculated with DESeq2 (where spike-in reads were used as control genes), filtered for non-standard chromosomes, then joined with the Unix function cat, sorted with the sort function (parameters:-k1,1-k2,2n), compressed with bgzip and indexed with tabix (parameters:-p bed) to be visualized within Washington University (WashU) Epigenome Browser (available online at http://epigenomegateway.wustl.edu/browser/)^91^. Tracks in BIGWIG format for ATAC-seq, CUT&RUN, and ChIP-seq data were generated with the DeepTools function bamCoverage (parameters:--normalizeUsing CPM--effectiveGenomeSize 2864785220 human or 2652783500 mouse--binSize 10--extendReads 300--ignoreDuplicates). The DeepTools function bigwigAverage was used to generate an average signal profile of ATAC-seq and CUT&RUN for each condition, while also filtering out signal from blacklist regions. Tracks were visualized either within the WashU epigenome browser or in tornado and aggregate signal plots generated with the DeepTools functions computeMatrix scale-regions (parameters:--startLabel Start--endLabel End--beforeRegionStartLength 3000--afterRegionStartLength 3000--skipZeros--missingDataAsZero); computeMatrix reference-point (parameters:--referencePoint center--beforeRegionStartLength 3000--afterRegionStartLength 3000-- skipZeros--missingDataAsZero); plotHeatmap (parameters:--xAxisLabel “” --yAxisLabel “Coverage” --heatmapHeight 12 --yMin 0 --refPointLabel “peak center”) with -- sortUsingSamples 1 in the tornado plots where MyoD ChIP-seq was the first track; plotProfile (parameters: --yAxisLabel “Coverage” --yMin 0 --refPointLabel “peak center” --perGroup -- startLabel Start --endLabel End). Hi-C and HiChIP signal in validPairs format for Hi-C data from (Dall’Agnese et al., 2019) from HiCPro^92^, or hic.txt format from MAPS (for HiChIP data) was merged with the UNIX function cat to create a single matrix for each condition and converted to HIC format with the Juicer Tools (v1.14.08)^93^ function pre (parameters: hg19-r 50000,25000,10000,5000,1000). Data were visualized either within the WashU epigenome browser (in HiC format, 5Kb resolution, KR normalization for Hi-C matrices, and longrangeformat for HiChIP loops) or with the R package GENOVA (v1.0.1)^94^ functions for aggregate peak analysis (APA, parameters: dist_thres = c(200e3, Inf)) or aggregate region analysis (ARA), after reading interaction matrices with the function load_contacts (parameters: resolution=5000 or 10000, balancing=’KR’), and syncing matrices from different conditions with the function sync_indices.

### 2D super-enhancers calls and 3D super-enhancers calling strategy

2D super-enhancers (SE) were called from H3K27ac ChIP-seq data with the Python script ROSE_main.py (Whyte et al, 2013)^52^ (parameters:-g hg19-t 2000-s 12500), with H3K27ac ChIP-seq Macs2 called peaks as input, and input chromatin as control. ROSE-derived SE were used as starting point for the calculation of 3D SE. For each experimental condition, ROSE-derived SEs were overlapped with the bins (at 5kb resolution) involved in H3K27ac HiChIP interactions. Bins belonging to those interactions were clustered with the BEDtools cluster function, specifying a max clustering distance of 10 kb (parameter:-d 10000), thus obtaining 3D SE. Within the 3D SE, hub enhancers were called according to (Huang et al., 2018)^55^. To further refine hub calls, hub bins were overlapped with H3K27ac ChIP-seq peaks with the BEDtools function intersect, to narrow down on 3D SE regulatory hotspots.

### Data integration

Overlaps of different genomic intervals was performed with the BEDtools function intersect, to retain a unique list of overlapping features (parameters:-wa-u), or to integrate different features to be processed within R for quantification (parameters:-wao), or to select only non-overlapping features (parameters:-v). Overlap of 3D super-enhancers was also filtered based on % of overlap between features (parameter:-f 0.6). Promoter regions were defined as-1000/+200 bp from the gene TSS, according to GENCODE.v19.annotation.gff3. Chromatin states were assigned to MyoD peaks considering their overlap with MyoD peaks summit. Partition of DARs chromatin states into genomic features (intergenic or intragenic: upstream, 5’-UTR, Exon, Intron, 3’-UTR, Downstream) was carried out according to PAVIS (available online at https://manticore.niehs.nih.gov/pavis2/)^95^.

### Re-analysis of public MuSCs data (RNA-seq, ATAC-seq, ChIP-seq)

Raw data in FASTQ format were retrieved from the Sequence Read Archive (SRA, https://www.ncbi.nlm.nih.gov/sra) at the following accession numbers: SRR16973998 to SRR16974007 for muSCs RNA-seq (4, 8, 16, 32, 60 hours post injury, 2 biological replicates for each condition), SRR16967713 to SRR16967722 for MuSCs ATAC-seq (same timepoints as RNA-seq), were previously described in Dong et al 2022^34^; SRR1200717 to SRR1200720 (*MyoD* ChIP-seq from primary myoblasts kept in growth media for three days, 2 biological replicates and their IgG controls) were previously described in Umansky et al 2015^96^. RNA- seq data were aligned to the mm10 Ensembl (November 2019) version of the mouse genome with STAR (parameters:--runMode alignReads--readFilesCommand zcat--outSAMtype BAM SortedByCoordinate--quantMode GeneCounts). Counts data (from STAR output in column 2, based on library preparation strandedness: unstranded) from all conditions were filtered based on their raw count, keeping genes where the sum of the counts for all samples was higher than 10. Differential expression analysis (using 4hpi as control condition) and Gene Ontology were carried out following the same workflow/thresholds as for our RNA-seq data. ATAC-seq and ChIP-seq data were aligned the mm10 Ensembl (November 2019) version of the mouse genome with Bowtie2 and followed the same analysis strategy as our ATAC-seq and ChIP-seq data.

## Data Availability

Sequencing data generated for this study have been deposited in the GEO database. Hi-C, H3K27ac ChIP-seq (IMR90), and MyoD ChIP-seq (IMR90/MyoD) can be found on GEO (GSE98530 and GSE128527). Public ATAC-seq and RNA-seq data from MuSCs can be found on GEO (GSE189074), as well as MyoD ChIP-seq in primary myoblasts (GSE56077).

## Acknowledgements

This work was supported by research grants from NIH, R01 GM134712-01 and R01 AR056712 to PLP; MDA Development Grant to LC (MDA 953791); CIRM postdoctoral training grant to CN; CIRM predoctoral training grant to MN; NIH NIGMS grant and NIH R01GM137117 to MDS; NIH R01 AR045203 to SJT; the Hong Kong Research Grant Council (AoE/M-604/16 and T13-605/18W) to THC. This study was partly supported by the Innovation and Technology Commission of Hong Kong (ITCPD/17-9) to THC. This publication includes data generated at the UC San Diego IGM Genomics Center utilizing an Illumina NovaSeq 6000 and Illumina NovaSeq X Plus that were purchased with funding from a National Institutes of Health SIG grant (#S10 OD026929)

