## Extended Figures for "E-box independent chromatin recruitment turns MYOD into a transcriptional repressor"

[1]Sanford Burnham Prebys Medical Discovery Institute, Development, Aging and Regeneration Program, La Jolla, CA. [2]Department of Molecular Biophysics & Biochemistry, Yale University, New Haven, CT. [3]Institute of Biomolecular Design & Discovery, Yale University, West Haven, CT. [4]Arima Genomics, San Diego, CA. [5]Division of Life Science, Center for Stem Cell Research, HKUST-Nan Fung Life Sciences Joint Laboratory, State Key Laboratory of Molecular Neuroscience, Daniel and Mayce Yu Molecular Neuroscience Center, The Hong Kong University of Science and Technology, Hong Kong, China. [6]Fred Hutchinson Cancer Center, Human Biology Division, Seattle, WA.

[#]These authors contributed equally to this manuscript

[‡] Current address: Department of Biomedical Engineering, Duke University, Durham, NC, USA

#### Extended Figure Legends

**Extended Fig. 1**

a) Schematic representation of human embryonic fibroblasts (IMR90) trans-differentiation in myoblasts through the ectopic expression of MYOD1. Within 24hrs, fibroblasts that were expressing GATA6 but not MYOD1 or MyHC turn off GATA6 and turn on myogenic lineage genes.
b) Representative image of immunofluorence analysis in IMR90 and IMR90/MyoD for MyoD (purple) and a myogenic lineage gene, MyHC (green).
c) Representative image of immunofluorence analysis in IMR90 and IMR90/MyoD for MyoD (magenta) and a fibroblast lineage gene, GATA6 (green).
d) qPCR of expression of myogenic lineage (*MYOD, ITGA7, TNNT2* and *MYH3*) and fibroblast lineage (*GATA6, FN1, IL6*), proliferation marker (*cFOS*) genes, with IMR90 levels in orange and IMR90/MyoD levels in blue.
e) Violin plot with the quantification of the IF signal in (c), with IMR90 in orange and IMR90/MyoD in blue.
n=3 biological replicates. Unpaired t-test

**Extended Fig. 2**

a-b) Gene Ontology of biological processes (upper panels) and cell type (lower panels) enriched in upregulated (a) or downregulated (b) genes in IMR90/MYOD vs IMR90 cells
c-d) Top10 enriched motifs for ATAC peaks at promoter (top) regions, with increased (c) or decreased (d) accessibility in IMR90/MYOD vs IMR90 cells

**Extended Fig. 3**

a) schematic representation of the binning of ATAC peaks in 3 regions
b-c) Aggregate signal of MyoD ChIP-seq (red), ATAC signal in IMR90 (orange) and IMR90/MyoD (blue) for DARs with increased (b) or decreased accessibility (c)
d-e) Top5 enriched motifs for ATAC peaks at promoter (top) regions, with increased (d) or decreased (e) accessibility in IMR90/MyoD vs IMR90 cells for genomic windows upstream of the summit (green), at the summit (yellow), and downstream of the summit (purple).

**Extended Fig. 4**

a-b) Tornado (left) and aggregate signal (right) plots for DARs with increased (a) or decreased (b) accessibility at differential genes promoters, showing MyoD CUT&RUN signal (orange), and the following (left to right) histone modifications in IMR90 and IMR90/MyoD: H3K27ac (blue), H3K4me3 (green), H3K4me1 (purple), H3K27me3 (red), H3K9me3 (grey).

c-d) Tornado (left) and aggregate signal (right) plots for DARs with increased (c) or decreased (d) accessibility at non-promoter regions, showing MyoD CUT&RUN signal (orange), and the following (left to right) histone modifications in IMR90 and IMR90/MyoD: H3K27ac (blue), H3K4me3 (green), H3K4me1 (purple), H3K27me3 (red), H3K9me3 (grey).

e) Tornado plots of regions that change chromatin state from active enhancer to active promoter, showing MyoD CUT&RUN signal (orange), and the following (left to right) histone modifications in IMR90 and IMR90/MyoD: H3K27ac (blue), H3K4me3 (green), H3K4me1 (purple), H3K27me3 (red), H3K9me3 (grey).

**Extended Fig. 5**

a-b) Aggregate signal (a) and tornado plots (b) of MyoD ChIP-seq signal and ATAC-seq signal for UP (a and b left panel) or DOWN (a and b right panel) DARs at non-promoter regions in IMR90/MyoD vs IMR90.

c-d) Aggregate signal (c) and tornado plots (d) of MyoD ChIP-seq signal and H3K27ac ChIP-seq signal for UP (c and d left panel) or DOWN (c and d right panel) DARs at enhancer candidates in IMR90/MyoD vs IMR90.

**Extended Fig. 6**

a) Aggregate signal of MyoD ChIP-seq and ATAC-seq signal at MyoD-bound non-promoter regions with increased (left, orange) or decreased (right, blue) chromatin accessibility in IMR90 vs IMR90/MyoD, and relative top5 chromatin states changes (lower panels); Q stands for quiescent state

b) Venn Diagram of super-enhancers (SE) in IMR90 and IMR90/MyoD, and those bound by MyoD.

c) Bar plot of number of MyoD peaks overlapping SE in IMR90 (blue) and IMR90/MyoD (green). Percentage represents fractions of total SE for that condition bound by MyoD.

d) Motifs enriched in MyoD peaks bound to SE present in IMR90 but lost in IMR90/MyoD (top panel), and SE present only in IMR90/MyoD (bottom panel).

e) Heatmap of Aggregate Peak Analysis (APA) of HiC data centered at conserved SE in IMR90 (left), IMR90/MyoD in growth condition (middle) and IMR90/MyoD in differentiation condition (right).

**Extended Fig. 7**

a-b) IGV screenshots of GATA6 (a) and KLF4 (b). Tracks from top to bottom: refseq gene; MYOD ChIP-seq; CTCF ChIP-seq in IMR90 (blue) and in IMR90/MYOD(orange); RNA-seq tracks in IMR90 (blue) and IMR90/MYOD (orange); ATAC-seq tracks in IMR90 (blue) and in IMR90/MYOD (orange); CUT&RUN for H3K27ac, H3K4me1, H3K4me3,H3K27me3, H3K9me3 in IMR90 (blue) and in IMR90/MYOD (orange); 3D-Superenhancer region in IMR90, H3K27ac HiChIP loops in IMR90 (blue), 3D-Superenhancer region in IMR90/Myod, H3K27ac HiChIP loops in IMR90/MyoD (orange); Hi-C connectivity in IMR90 (blue) and in IMR90/MYOD (orange).

c) Barplot of % of CTCF differential HiChIP interaction overlapping 3D-SEs either lost or gained in IMR90/MyoD vs IMR90 whose interaction are increased or decreased.

d) Aggregate plots (top) and tornado plots (bottom) of MYOD ChIP-seq, ATAC-seq, Kacme-ChIP-seq and CTCF-ChIP-seq overlapping 3D SEs either lost or gained in IMR90/MyoD vs IMR90.

**Extended Fig. 8**

IGV screenshots of *IL1b*, *cFOS*, and *KLF4* (examples of repressed genes) and *MYOG* (example of activated gene) loci. Tracks from top to bottom: refseq gene; ATAC-seq; RNA-seq; MYOD CUT&RUN; Kacme CUT&RUN at 4hrs (blue) and 60hrs (orange); MYOD ChIP-seq in mouse primary myoblasts from Umansky et al 2015.

**Extended Fig. 9**

a) Venn Diagram of the overlap between genes in IMR90/MyoD vs IMR90 that are not repressed by various MYOD mutant construct.

b-g) Gene Ontology of biological processes of genes in (a).

Ext. Fig. 1

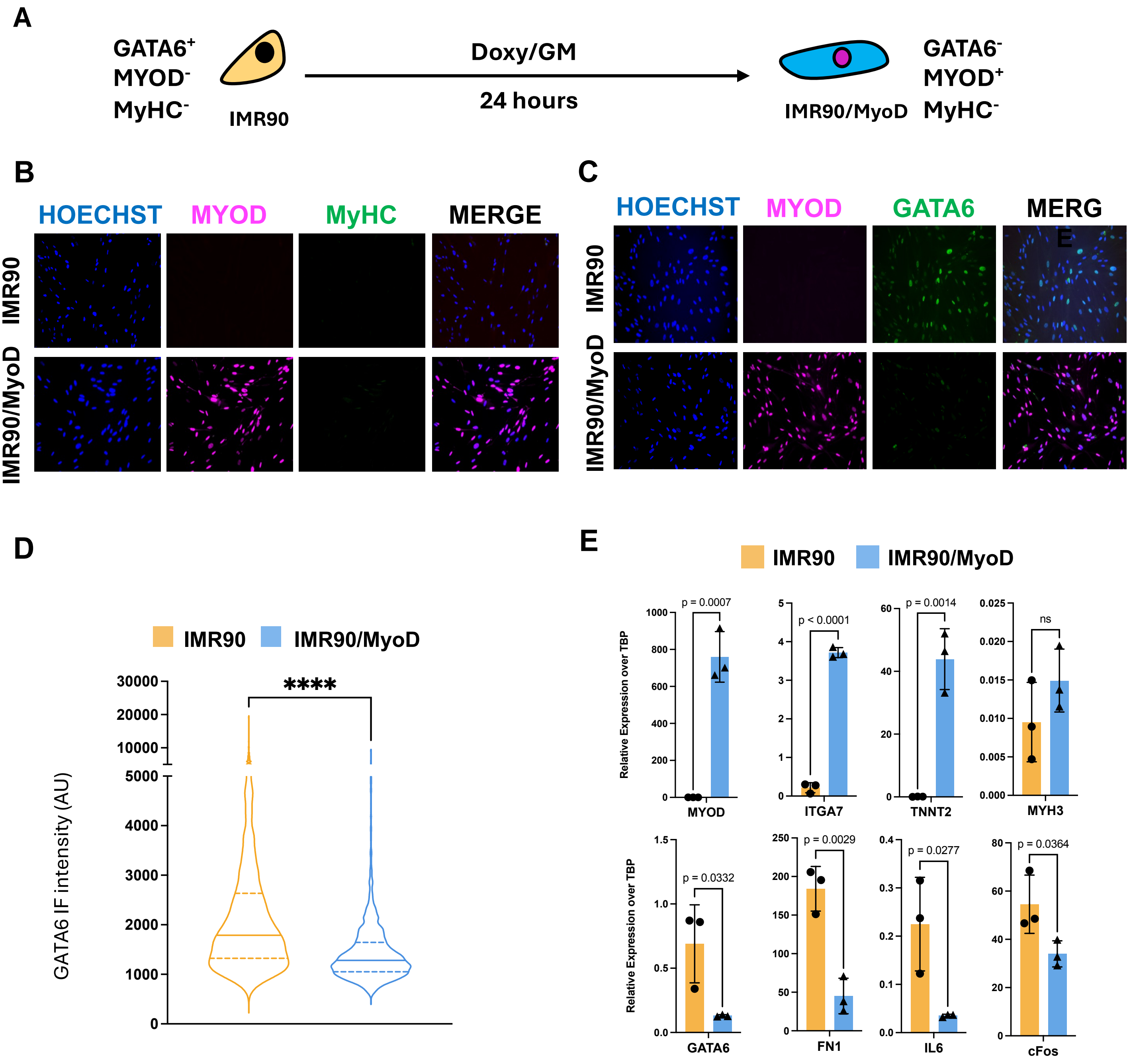

A
GATA6+
MYOD-
MyHC-
IMR90
Doxy/GM
24 hours
IMR90/MyoD
GATA6-
MYOD+
MyHC-
B
HOECHST
MYOD
MyHC
MERGE
IMR90
IMR90/MyoD
C
HOECHST
MYOD
GATA6
MERG
E
IMR90
IMR90/MyoD
D
IMR90
IMR90/MyoD
GATA6 IF intensity (AU)
****
E
IMR90
IMR90/MyoD
Relative Expression over TBP
p = 0.0007
MYOD
p < 0.0001
ITGA7
p = 0.0014
TNNT2
ns
MYH3
p = 0.0332
GATA6
p = 0.0029
FN1
p = 0.0277
IL6
p = 0.0364
cFos

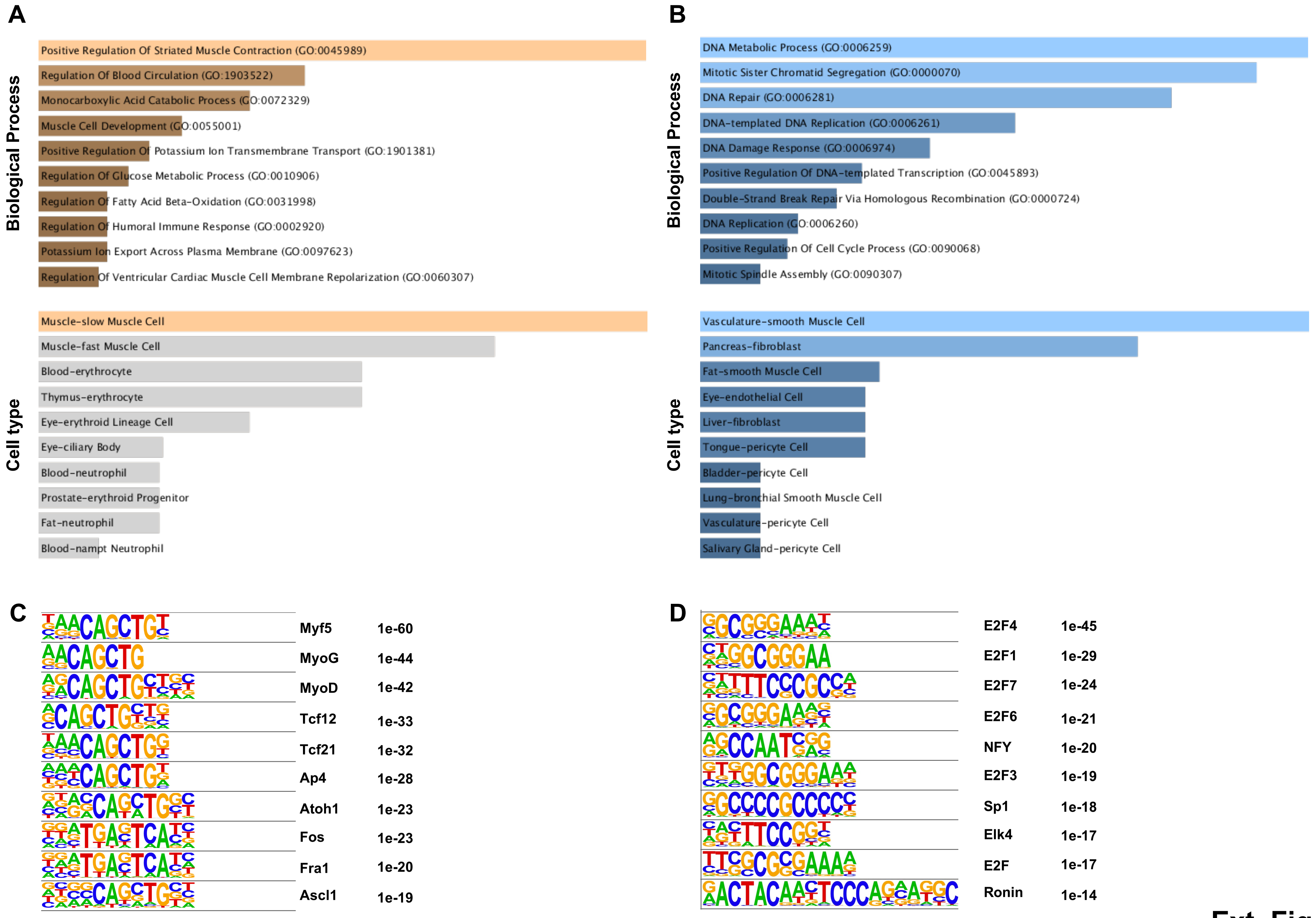
A
Biological Process
Positive Regulation Of Striated Muscle Contraction (GO:0045989)
Regulation Of Blood Circulation (GO:1903522)
Monocarboxylic Acid Catabolic Process (GO:0072329)
Muscle Cell Development (GO:0055001)
Positive Regulation Of Potassium Ion Transmembrane Transport (GO:1901381)
Regulation Of Glucose Metabolic Process (GO:0010906)
Regulation Of Fatty Acid Beta-Oxidation (GO:0031998)
Regulation Of Humoral Immune Response (GO:0002920)
Potassium Ion Export Across Plasma Membrane (GO:0097623)
Regulation Of Ventricular Cardiac Muscle Cell Membrane Repolarization (GO:0060307)
Cell type
Muscle-slow Muscle Cell
Muscle-fast Muscle Cell
Blood-erythrocyte
Thymus-erythrocyte
Eye-erythroid Lineage Cell
Eye-ciliary Body
Blood-neutrophil
Prostate-erythroid Progenitor
Fat-neutrophil
Blood-nampt Neutrophil
B
Biological Process
DNA Metabolic Process (GO:0006259)
Mitotic Sister Chromatid Segregation (GO:0000070)
DNA Repair (GO:0006281)
DNA-templated DNA Replication (GO:0006261)
DNA Damage Response (GO:0006974)
Positive Regulation Of DNA-templated Transcription (GO:0045893)
Double-Strand Break Repair Via Homologous Recombination (GO:0000724)
DNA Replication (GO:0006260)
Positive Regulation Of Cell Cycle Process (GO:0090068)
Mitotic Spindle Assembly (GO:0090307)
Cell type
Vasculature-smooth Muscle Cell
Pancreas-fibroblast
Fat-smooth Muscle Cell
Eye-endothelial Cell
Liver-fibroblast
Tongue-pericyte Cell
Bladder-pericyte Cell
Lung-bronchial Smooth Muscle Cell
Vasculature-pericyte Cell
Salivary Gland-pericyte Cell
C
Myf5 1e-60
MyoG 1e-44
MyoD 1e-42
Tcf12 1e-33
Tcf21 1e-32
Ap4 1e-28
Atoh1 1e-23
Fos 1e-23
Fra1 1e-20
Ascl1 1e-19
D
E2F4 1e-45
E2F1 1e-29
E2F7 1e-24
E2F6 1e-21
NFY 1e-20
E2F3 1e-19
Sp1 1e-18
Elk4 1e-17
E2F 1e-17
Ronin 1e-14

Ext. Fig. 2

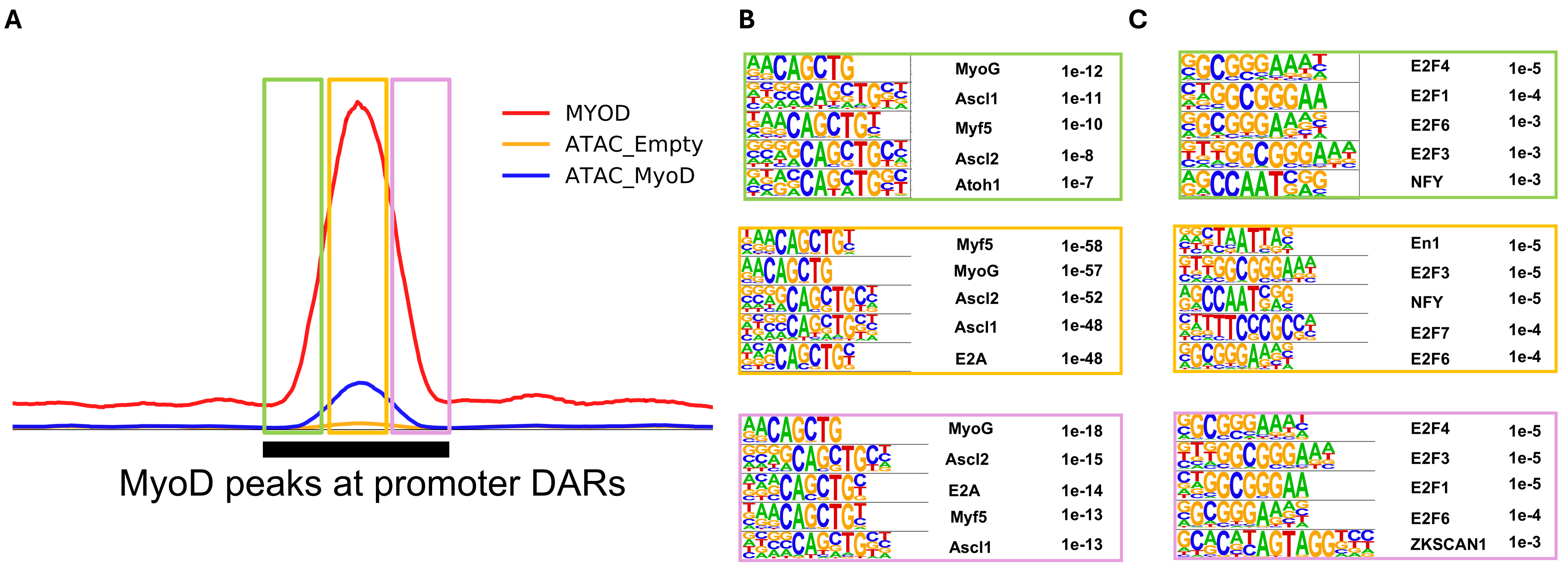

A
MYOD
ATAC_Empty
ATAC_MyoD
MyoD peaks at promoter DARs
B
MyoG 1e-12
Ascl1 1e-11
Myf5 1e-10
Ascl2 1e-8
Atoh1 1e-7
Myf5 1e-58
MyoG 1e-57
Ascl2 1e-52
Ascl1 1e-48
E2A 1e-48
MyoG 1e-18
Ascl2 1e-15
E2A 1e-14
Myf5 1e-13
Ascl1 1e-13
C
E2F4 1e-5
E2F1 1e-4
E2F6 1e-3
E2F3 1e-3
NFY 1e-3
En1 1e-5
E2F3 1e-5
NFY 1e-5
E2F7 1e-4
E2F6 1e-4
E2F4 1e-5
E2F3 1e-5
E2F1 1e-5
E2F6 1e-4
ZKSCAN1 1e-3


Ext. Fig. 3

### Ext Fig. 4

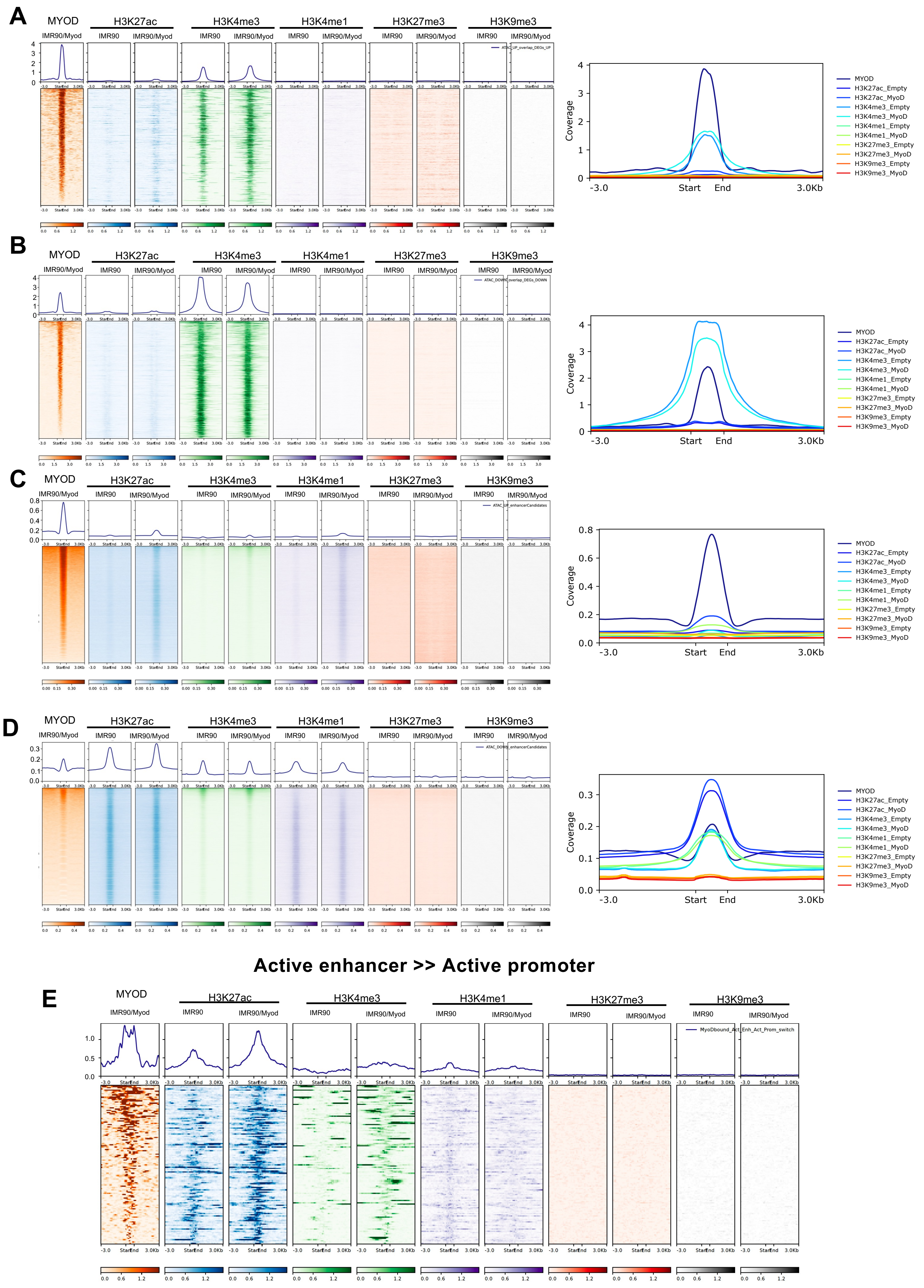
A
MYOD
IMR90/Myod
H3K27ac
H3K4me3
H3K4me1
H3K27me3
H3K9me3
IMR90
IMR90/Myod
ATAC_UP_overlap_DEGs_UP
Coverage
-3.0
Start
End
3.0Kb
MYOD
H3K27ac_Empty
H3K27ac_MyoD
H3K4me3_Empty
H3K4me3_MyoD
H3K4me1_Empty
H3K4me1_MyoD
H3K27me3_Empty
H3K27me3_MyoD
H3K9me3_Empty
H3K9me3_MyoD
B
ATAC_DOWN_overlap_DEGs_DOWN
C
ATAC_UP_enhancerCandidates
D
ATAC_DOWN_enhancerCandidates
Active enhancer >> Active promoter
E
MyoDbound_Act_Enh_Act_Prom_switch

Ext. Fig. 5

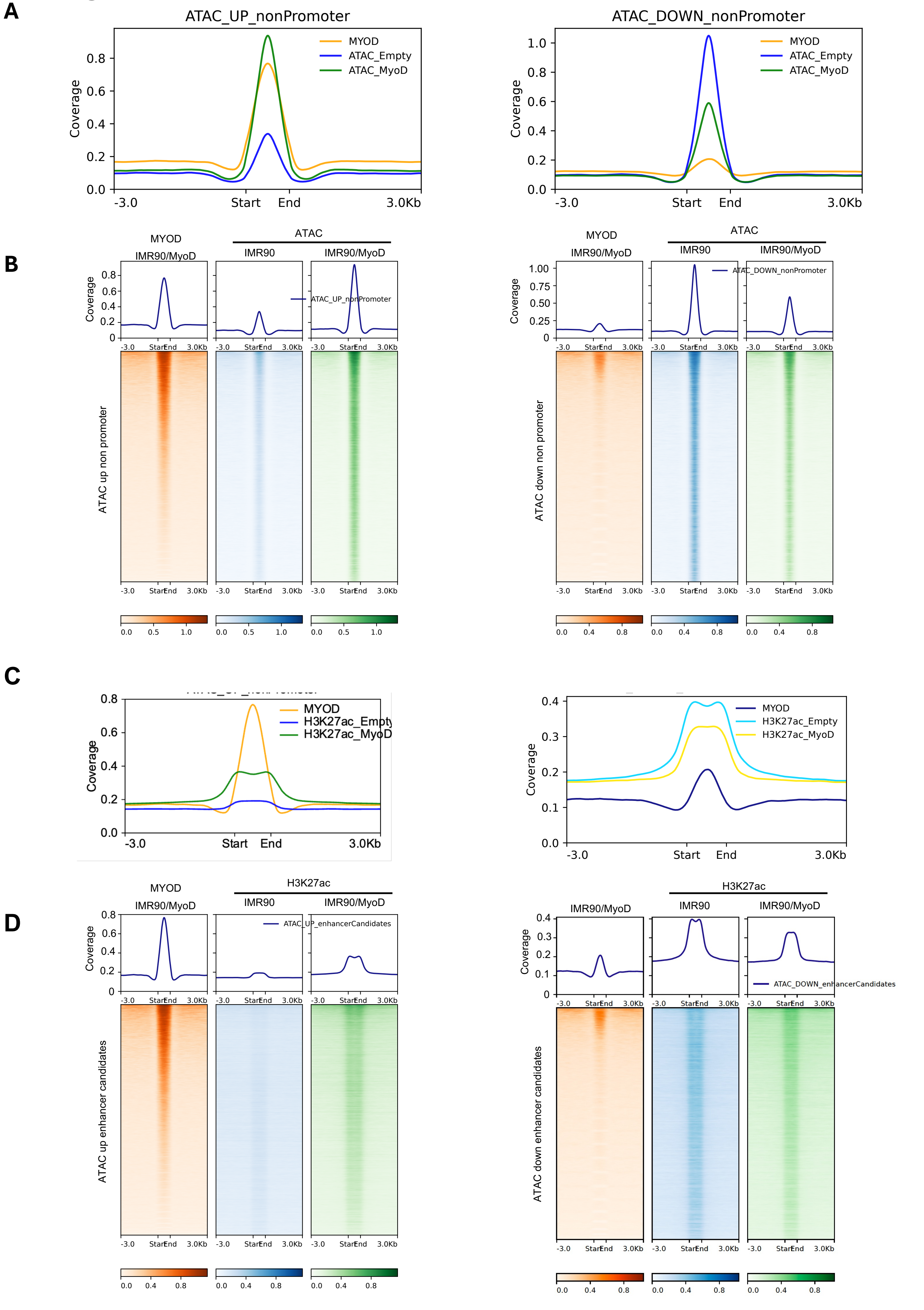
A
ATAC_UP_nonPromoter
MYOD
ATAC_Empty
ATAC_MyoD
Coverage
-3.0
Start
End
3.0Kb
ATAC_DOWN_nonPromoter
B
MYOD
IMR90/MyoD
ATAC
IMR90
ATAC_UP_nonPromoter
ATAC up non promoter
ATAC_DOWN_nonPromoter
ATAC down non promoter
C
H3K27ac_Empty
H3K27ac_MyoD
D
H3K27ac
ATAC_UP_enhancerCandidates
ATAC up enhancer candidates
ATAC_DOWN_enhancerCandidates
ATAC down enhancer candidates

### Ext. Fig. 6

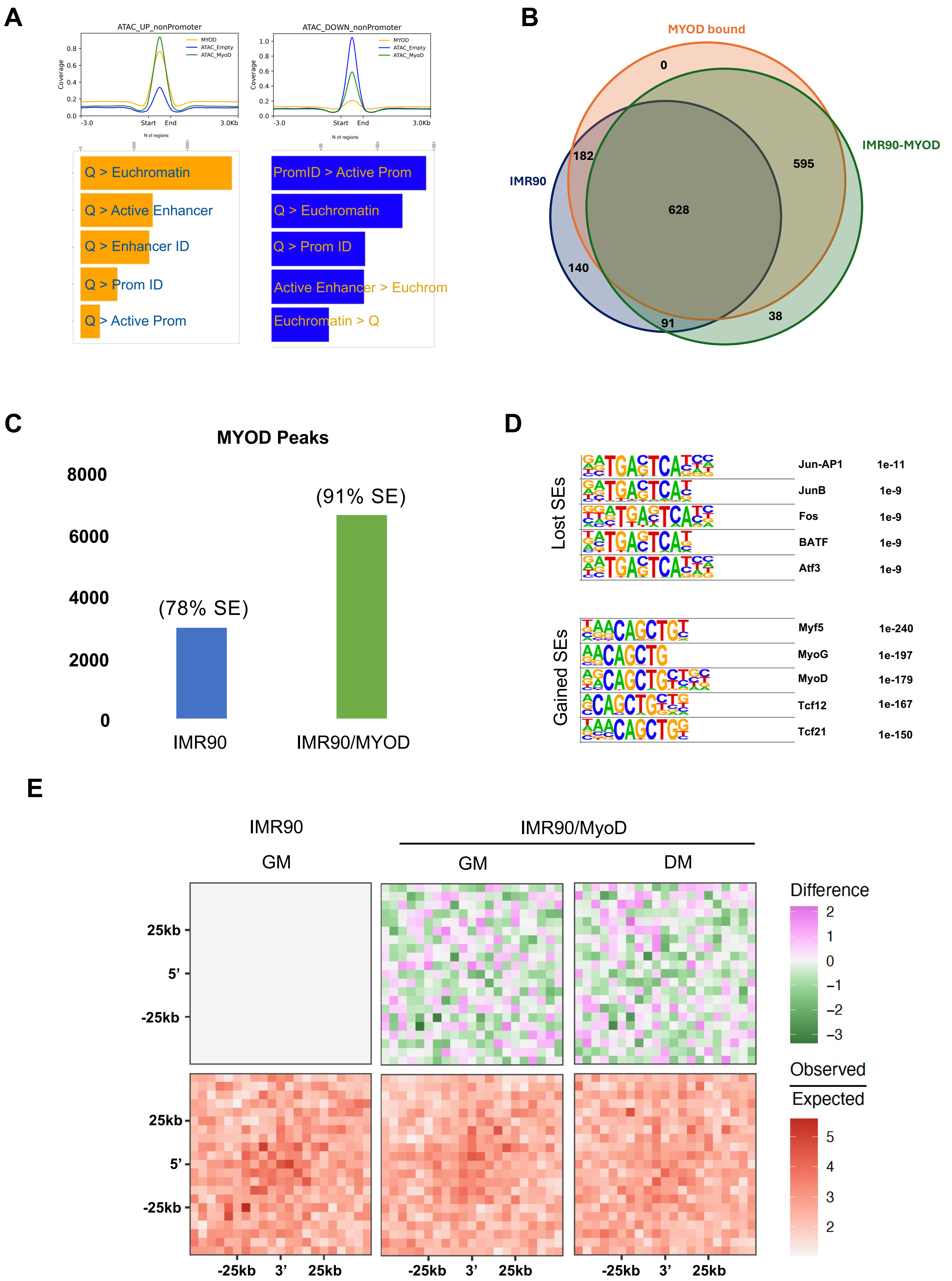
A
ATAC_UP_nonPromoter
MYOD
ATAC_Empty
ATAC_MyoD
Coverage
-3.0
Start
End
3.0Kb
N of regions
Q > Euchromatin
Q > Active Enhancer
Q > Enhancer ID
Q > Prom ID
Q > Active Prom
ATAC_DOWN_nonPromoter
PromID > Active Prom
Q > Euchromatin
Q > Prom ID
Active Enhancer > Euchrom
Euchromatin > Q
B
MYOD bound
IMR90-MYOD
IMR90
0
182
595
628
140
91
38
C
MYOD Peaks
8000
6000
4000
2000
0
(78% SE)
(91% SE)
IMR90
IMR90/MYOD
D
Lost SEs
Jun-AP1 1e-11
JunB 1e-9
Fos 1e-9
BATF 1e-9
Atf3 1e-9
Gained SEs
Myf5 1e-240
MyoG 1e-197
MyoD 1e-179
Tcf12 1e-167
Tcf21 1e-150
E
IMR90
IMR90/MyoD
GM
GM
DM
25kb
5'
-25kb
-25kb
3'
25kb
Difference
2
1
0
−1
−2
−3
Observed
Expected
5
4
3
2

### Ext. Figure 7

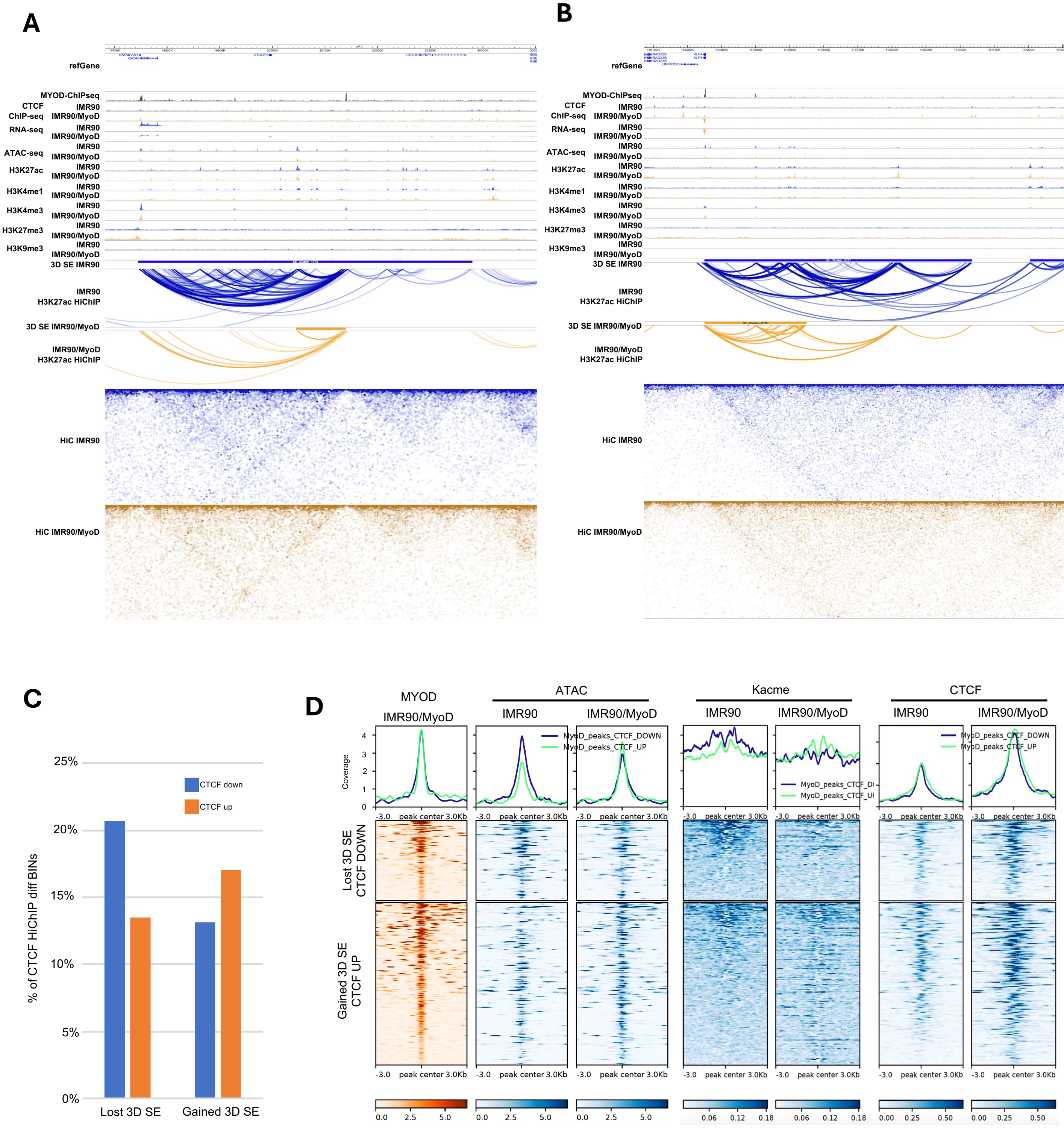

A
B
refGene
MYOD-ChIPseq
CTCF ChIP-seq
RNA-seq
ATAC-seq
H3K27ac
H3K4me1
H3K4me3
H3K27me3
H3K9me3
IMR90
IMR90/MyoD
3D SE IMR90
IMR90 H3K27ac HiChIP
3D SE IMR90/MyoD
IMR90/MyoD H3K27ac HiChIP
HiC IMR90
HiC IMR90/MyoD
C
CTCF down
CTCF up
% of CTCF HiChIP diff BINs
25%
20%
15%
10%
5%
0%
Lost 3D SE
Gained 3D SE
D
MYOD
ATAC
Kacme
CTCF
IMR90
IMR90/MyoD
Coverage
MyoD_peaks_CTCF_DOWN
MyoD_peaks_CTCF_UP
-3.0
peak center 3.0Kb
Lost 3D SE CTCF DOWN
Gained 3D SE CTCF UP
0.0 2.5 5.0
0.06 0.12 0.18
0.00 0.25 0.50

### Ext. Figure 8

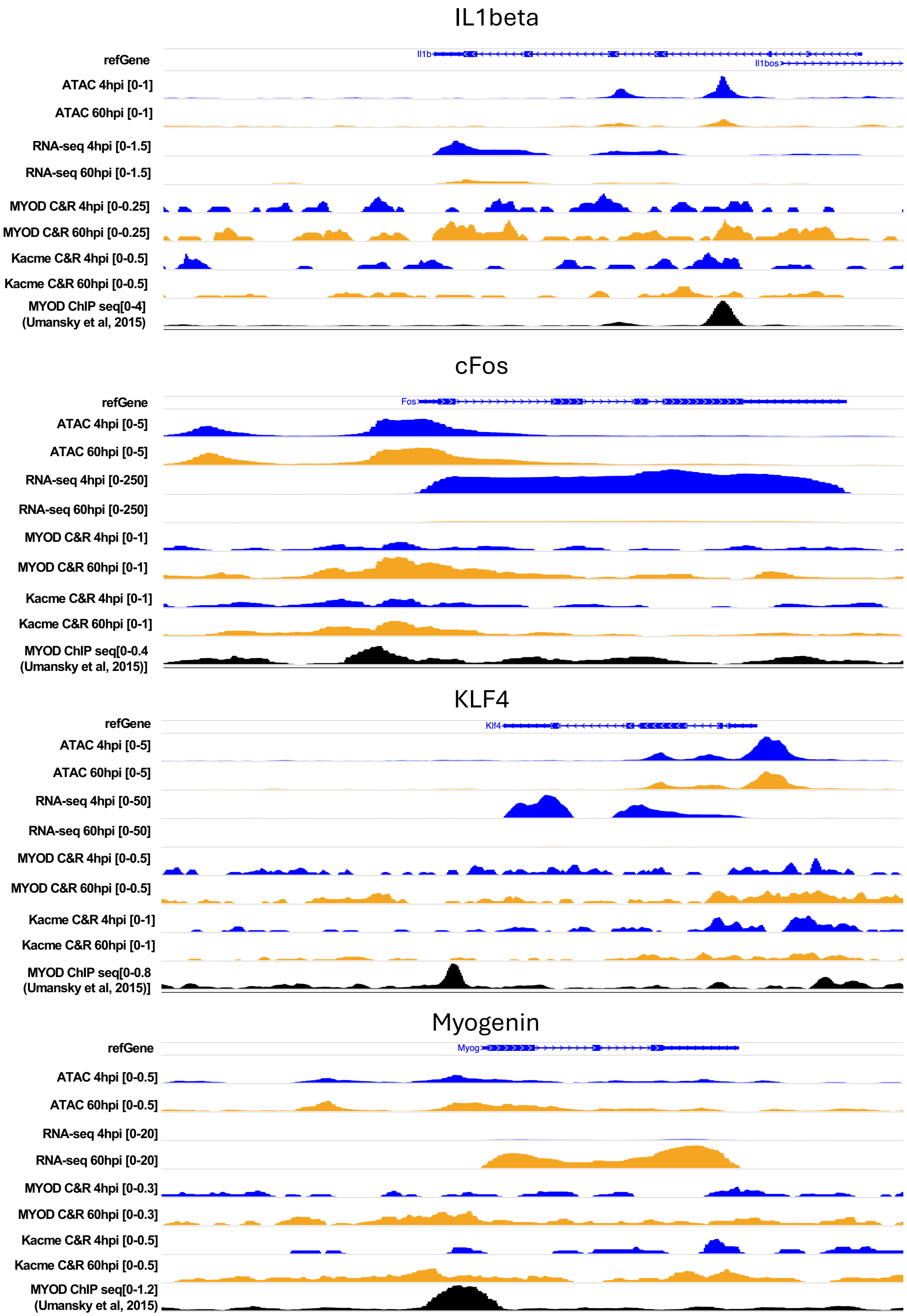

IL1beta
refGene
Il1b
Il1bos
ATAC 4hpi [0-1]
ATAC 60hpi [0-1]
RNA-seq 4hpi [0-1.5]
RNA-seq 60hpi [0-1.5]
MYOD C&R 4hpi [0-0.25]
MYOD C&R 60hpi [0-0.25]
Kacme C&R 4hpi [0-0.5]
Kacme C&R 60hpi [0-0.5]
MYOD ChIP seq[0-4]
(Umansky et al, 2015)
cFos
refGene
Fos
ATAC 4hpi [0-5]
ATAC 60hpi [0-5]
RNA-seq 4hpi [0-250]
RNA-seq 60hpi [0-250]
MYOD C&R 4hpi [0-1]
MYOD C&R 60hpi [0-1]
Kacme C&R 4hpi [0-1]
Kacme C&R 60hpi [0-1]
MYOD ChIP seq[0-0.4
(Umansky et al, 2015)]
KLF4
refGene
Klf4
ATAC 4hpi [0-5]
ATAC 60hpi [0-5]
RNA-seq 4hpi [0-50]
RNA-seq 60hpi [0-50]
MYOD C&R 4hpi [0-0.5]
MYOD C&R 60hpi [0-0.5]
Kacme C&R 4hpi [0-1]
Kacme C&R 60hpi [0-1]
MYOD ChIP seq[0-0.8
(Umansky et al, 2015)]
Myogenin
refGene
Myog
ATAC 4hpi [0-0.5]
ATAC 60hpi [0-0.5]
RNA-seq 4hpi [0-20]
RNA-seq 60hpi [0-20]
MYOD C&R 4hpi [0-0.3]
MYOD C&R 60hpi [0-0.3]
Kacme C&R 4hpi [0-0.5]
Kacme C&R 60hpi [0-0.5]
MYOD ChIP seq[0-1.2]
(Umansky et al, 2015)

### Ext. Figure 9

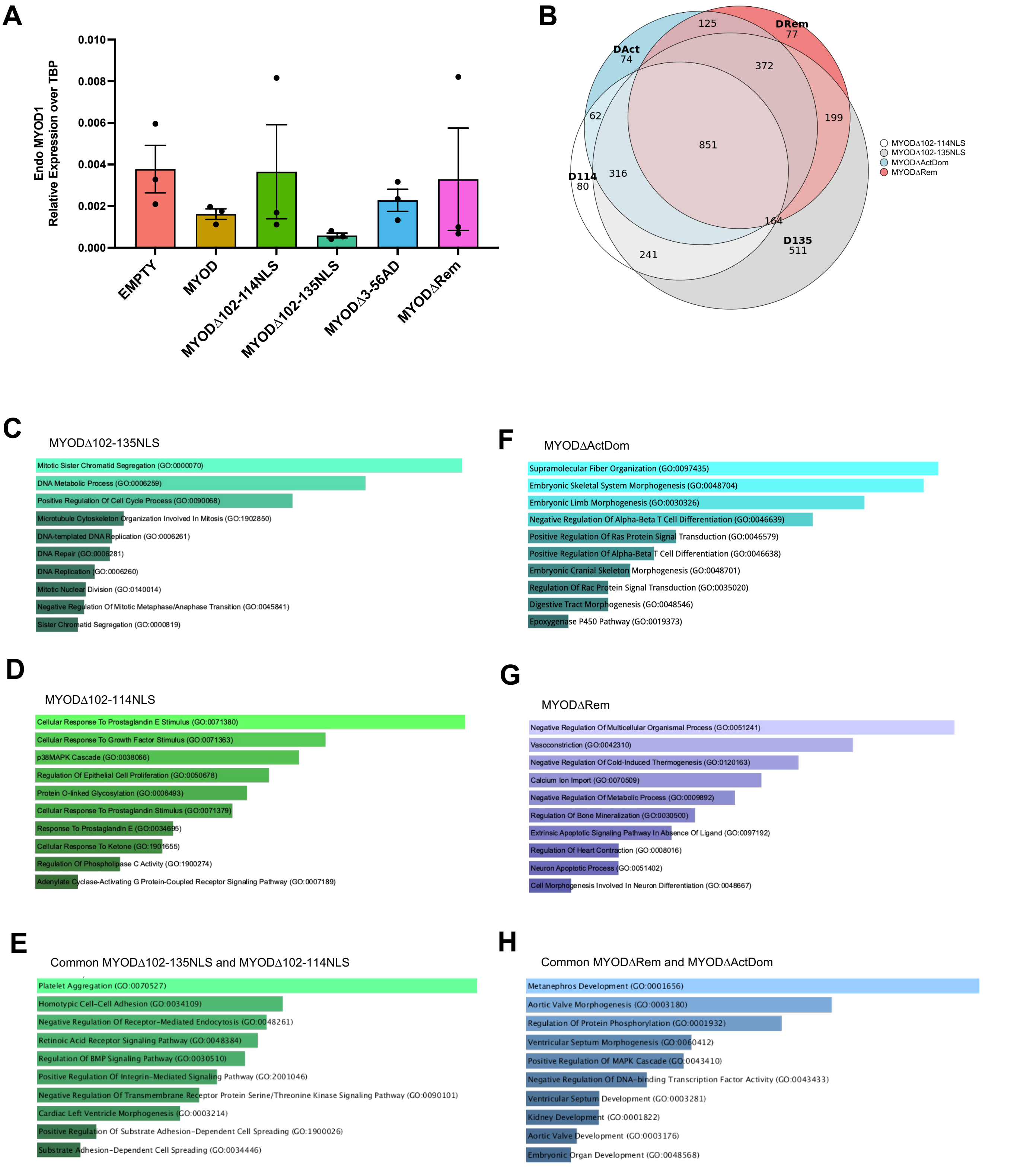
A
Endo MYOD1
Relative Expression over TBP
0.010
0.008
0.006
0.004
0.002
0.000
EMPTY
MYOD
MYODΔ102-114NLS
MYODΔ102-135NLS
MYODΔ3-56AD
MYODΔRem
B
125
DRem
77
DAct
74
372
62
199
851
D114
80
316
164
D135
511
241
MYODΔ102-114NLS
MYODΔ102-135NLS
MYODΔActDom
MYODΔRem
C
MYODΔ102-135NLS
Mitotic Sister Chromatid Segregation (GO:0000070)
DNA Metabolic Process (GO:0006259)
Positive Regulation Of Cell Cycle Process (GO:0090068)
Microtubule Cytoskeleton Organization Involved In Mitosis (GO:1902850)
DNA-templated DNA Replication (GO:0006261)
DNA Repair (GO:0006281)
DNA Replication (GO:0006260)
Mitotic Nuclear Division (GO:0140014)
Negative Regulation Of Mitotic Metaphase/Anaphase Transition (GO:0045841)
Sister Chromatid Segregation (GO:0000819)
F
MYODΔActDom
Supramolecular Fiber Organization (GO:0097435)
Embryonic Skeletal System Morphogenesis (GO:0048704)
Embryonic Limb Morphogenesis (GO:0030326)
Negative Regulation Of Alpha-Beta T Cell Differentiation (GO:0046639)
Positive Regulation Of Ras Protein Signal Transduction (GO:0046579)
Positive Regulation Of Alpha-Beta T Cell Differentiation (GO:0046638)
Embryonic Cranial Skeleton Morphogenesis (GO:0048701)
Regulation Of Rac Protein Signal Transduction (GO:0035020)
Digestive Tract Morphogenesis (GO:0048546)
Epoxygenase P450 Pathway (GO:0019373)
D
MYODΔ102-114NLS
Cellular Response To Prostaglandin E Stimulus (GO:0071380)
Cellular Response To Growth Factor Stimulus (GO:0071363)
p38MAPK Cascade (GO:0038066)
Regulation Of Epithelial Cell Proliferation (GO:0050678)
Protein O-linked Glycosylation (GO:0006493)
Cellular Response To Prostaglandin Stimulus (GO:0071379)
Response To Prostaglandin E (GO:0034695)
Cellular Response To Ketone (GO:1901655)
Regulation Of Phospholipase C Activity (GO:1900274)
Adenylate Cyclase-Activating G Protein-Coupled Receptor Signaling Pathway (GO:0007189)
G
MYODΔRem
Negative Regulation Of Multicellular Organismal Process (GO:0051241)
Vasoconstriction (GO:0042310)
Negative Regulation Of Cold-Induced Thermogenesis (GO:0120163)
Calcium Ion Import (GO:0070509)
Negative Regulation Of Metabolic Process (GO:0009892)
Regulation Of Bone Mineralization (GO:0030500)
Extrinsic Apoptotic Signaling Pathway In Absence Of Ligand (GO:0097192)
Regulation Of Heart Contraction (GO:0008016)
Neuron Apoptotic Process (GO:0051402)
Cell Morphogenesis Involved In Neuron Differentiation (GO:0048667)
E
Common MYODΔ102-135NLS and MYODΔ102-114NLS
Platelet Aggregation (GO:0070527)
Homotypic Cell-Cell Adhesion (GO:0034109)
Negative Regulation Of Receptor-Mediated Endocytosis (GO:0048261)
Retinoic Acid Receptor Signaling Pathway (GO:0048384)
Regulation Of BMP Signaling Pathway (GO:0030510)
Positive Regulation Of Integrin-Mediated Signaling Pathway (GO:2001046)
Negative Regulation Of Transmembrane Receptor Protein Serine/Threonine Kinase Signaling Pathway (GO:0090101)
Cardiac Left Ventricle Morphogenesis (GO:0003214)
Positive Regulation Of Substrate Adhesion-Dependent Cell Spreading (GO:1900026)
Substrate Adhesion-Dependent Cell Spreading (GO:0034446)
H
Common MYODΔRem and MYODΔActDom
Metanephros Development (GO:0001656)
Aortic Valve Morphogenesis (GO:0003180)
Regulation Of Protein Phosphorylation (GO:0001932)
Ventricular Septum Morphogenesis (GO:0060412)
Positive Regulation Of MAPK Cascade (GO:0043410)
Negative Regulation Of DNA-binding Transcription Factor Activity (GO:0043433)
Ventricular Septum Development (GO:0003281)
Kidney Development (GO:0001822)
Aortic Valve Development (GO:0003176)
Embryonic Organ Development (GO:0048568)
